# Human DDIT4L intron retention contributes to cognitive impairment and amyloid plaque formation

**DOI:** 10.1101/2023.12.30.573740

**Authors:** Kai-Cheng Li, Hai-Xiang Shi, Zhen Li, Pu You, Jing Pan, Yi-Chuan Cai, Jin-Wen Li, Xue-Fei Ma, Shuo Zhang, Lei Diao, Bing Cai, Yang Lu, Hai-Bo Wang, Yan-Qing Zhong, Liang Chen, Ying Mao, Xu Zhang

## Abstract

Cognitive impairment and amyloid plaques are the most important clinical and neuropathological feature for dementia, especially in Alzheimer’s disease (AD). However, the etiology of dementia is complicated. The present study reveals that an aberrant splicing of DDIT4L, the isoform DDIT4L intron retention (DIR), occurs in AD patients. Homozygous DIR-knock-in (KI) mice showed DIR expression in hippocampal neurons, marked cognitive impairment, augmented Aβ deposition and enhanced Tau phosphorylation. The DIR colocalized with thioflavin S-positive plaques and gelsolin in AD patients. The DIR induced Aβ deposition and cognitive impairment by interacting with gelsolin. Moreover, DIR interacted with GluA1, the subunit of the AMPA receptor, contributing to synaptic deficiency and cognitive impairment. Furthermore, an anti-DIR monoclonal antibody (mAb) alleviated cognitive impairment and reduced Aβ deposition and Tau phosphorylation. Thus, DIR contributes to cognitive impairment and amyloid plaques, and could be a potential therapeutic target for dementia.

## Introduction

Alzheimer’s disease (AD) is the most common form of age-related dementia^1,2^. The clinical manifestations of AD include profound cognitive decline, progressive memory loss, and retrograde and anterograde amnesia, which are accompanied by severe histopathological changes, such as degeneration of the hippocampus and subsequent loss of cortical matter^3^. Histopathological changes in AD patients can be observed decades earlier than the onset of clinical manifestations^4^. Current opinions suggest that the aggregation of amyloid β (Aβ), followed by tau accumulation, occurs in the brain prior to the onset of cognitive defects^5,6^. Many studies have shown that Aβ peptides are prone to aggregate into β sheet conformations, which is the main form of dense-core plaques, and these aggregates are observed in the brains of AD patients^7-10^. Interestingly, Aβ peptides that are extracted from the brains of AD patients, but not that from the brains of control subjects without AD, can act as a seed for fibril formation *in vitro*^11,12^. Besides certain fibrillar structures of Aβ, the explanation for these findings is that other factors in pathological plaques have amplified amyloid deposition^13,14^. While whether have unidentified factors inducing plaque formation? On the other hand, previous studies showed that Aβ, especially Aβ oligomers, decreased synaptic transmission via postsynaptic NMDA receptors, metabotropic glutamate receptor 5 and α7 nicotinic acetylcholine receptor^15-17^, accounting for cognitive impairment in AD animal models. Ongoing efforts in suppressing Aβ formation or eliminating toxic Aβ have a modest clinical effect^18^. These data suggest that there may be unidentified factors, besides Aβ, also contributing to cognitive impairment in dementia patients.

Cerebral hypoxia is believed to be a risk factor for AD because of its roles in cognitive impairment, Aβ accumulation and enhanced Tau phosphorylation^19-22^. Cerebral hypoxia can be caused by several diseases, such as cerebral vascular diseases, cardiovascular diseases, respiratory disorders, cervical spondylopathy and anemia. Brain tumors may also cause local hypoxia due to insufficient blood supply^23^. Recently, hypoxia was linked to Aβ deposition in young COVID patients and found to be associated with AD via induction of sustained neuroinflammation in the brain microenvironment^24^. Hypoxia was also reported to influence mRNA splicing^25,26^. Alternative splicing (AS) of pre-mRNA transcripts is a fundamental regulatory process that increases proteome diversity^27^. One form of AS, intron retention (IR) of pre-mRNAs, may contribute to AD development or be a target for AD treatment^28-31^. The present study shows that hypoxia-induced aberrant splicing of human DNA damage inducible transcript 4 like (DDIT4L), resulting in the production of an IR isoform (named as DDIT4L intron retention (DIR)), occurs in AD patients. Homozygous DIR-knock-in (KI) mice displayed cognitive decline and exhibited the major pathological characteristics of AD. Interestingly, DIR contributed Aβ deposition, plaque formation and cognitive impairment by interacting with the C228-N233 domain of gelsolin and DIR bound the R198-E205 domain in the GluA1 subunit of the AMPA receptor, which mediated DIR-induced synaptic deficiency and cognitive impairment. A monoclonal antibody (mAb) against DIR relieved cognitive impairment and reduced Aβ deposition and Tau phosphorylation in the DIR-KI mice.

## Results

### Aberrant splicing of human DDIT4L in AD patients

In our previous study, to identify potential candidates for predicting the prognosis of glioblastoma (GBM), we examined upregulated genes in GBM and low-grade glioma (LGG) samples with data from the TCGA database. We found that human DDIT4L was one of the genes that was most highly upregulated in GBM (Supplementary Fig. S1a and unpublicated data for DDIT4L in glioma). During the experiment, PCR analysis of human DDIT4L mRNA expression in GBM tissues unexpectedly revealed that in addition to the predicted ∼200 bp product, there was an extra ∼2000 bp product (Fig. 1a). Then in this study, we focused on the sequence for the longer product and found that it was DIR (Fig. 1a). The human *DDIT4L* gene consists of three exons with a start codon in exon 2 and encodes a protein of 193 amino acids (AAs). *DIR* shares exon 2 with DDIT4L but has a unique intron, which translates to 84 AAs (Fig. 1b; Supplementary Fig. S4a). We prepared an antibody against the 30 AAs of the translated sequence of exon 2 and tested it in patient tissues. Immunoblotting of GBM tissues revealed two bands (∼12 kDa and ∼22 kDa) (Fig. 1c). Since the molecular weight of DDIT4L is ∼22 kDa, we speculated that the 12 kDa band might be DIR. Then, we generated an antibody that recognized only the 54 AAs of the translated sequence of the intron unique to DIR and detected an ∼12 kDa protein in the GBM tissues (Fig. 1d). Furthermore, we constructed a heterologous expression plasmid containing the open reading frame of *DIR* and transfected the plasmid into HEK293 cells. DIR could be detected in the cell lysates as well as the conditioned medium (Fig. 1e), indicating that *DIR* could be translated and released *in vivo*.

**Fig. 1:**
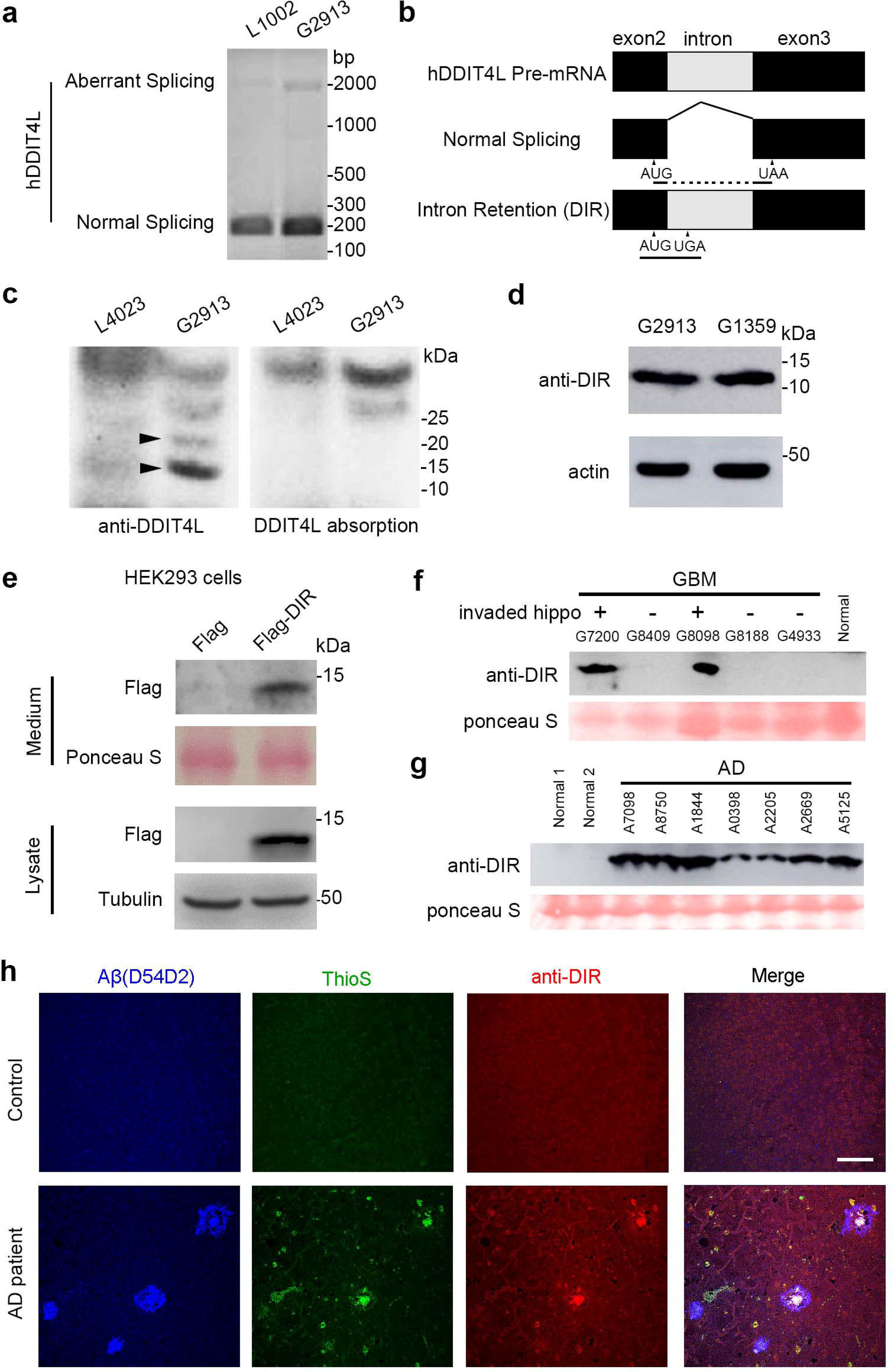
DIR is an aberrant splicing of human DDIT4L in the patients. a. There were two products (∼200 bp and ∼2000 bp) revealed by the PCR experiment with the primers for human DDIT4L mRNA prepared from the GBM tissue of patient 1 (G2913), but not in low grade glioma (LGG) tissue from the LGG patient 1 (L1002). b. The DIR was a splicing isoform of human DDIT4L with the retention of intron between the exon 2 and exon 3. c. Immunoblotting of the GBM extract showed two immunoreactive bands (∼22 kDa and ∼12 kDa, arrowheads) detected with the antibody against the N-terminal of DDIT4L, but not by the pre-absorbed antibody. d. In the GBM extract from two patient (G2913, G1359), ∼12 kDa immunoblot was detected with the DIR antibody against the intron translated region. e. In the HEK293 cells transfected with the plasmid expressing Flag-DIR, the Flag-DIR could be detected in both the cell lysate and the culture medium. f. The DIR was mainly detected in the plasma from the patients with GBM invaded the hippocampus (G8098 and G7200). g. The DIR was found in the AD patients’ plasma, named as A7098, etc. h. In AD patients (n = 6), but not control individuals (n = 6), Aβ- and thioflavine S (ThioS)-positive dense-core plaque co-localized with DIR in the hippocampal DG area. Scale bar = 50 mm.

We also produced antibodies targeting the last 27 AAs of DIR to test whether DIR existed in the blood of GBM patients. Immunoblotting showed that high levels of DIR were present in the blood of patients with tumors invading the hippocampus (Fig. 1f; Supplementary Fig. S1b, S2c). Statistics showed that DIR was present in blood samples from 71.4% of the patients with GBM invading the hippocampus (n = 7) but the proportion was only 15.4% of patients with GBM not invading the hippocampus (n = 13), suggesting a high correlation between DIR production and tumor invasion of the hippocampus, which is important for the storage of declarative memory and associative learning.

On the other hand, GBM always induces hypoxic conditions in solid tumor tissues^23,32^. For DDIT4L could be upregulated by hypoxia (Fig. S7a), we also tested whether hypoxia induced the expression of *DIR* in neurons. The *DIR* protein expression was increased in neurons cultured under hypoxic conditions compared with control neurons (Supplementary Fig. S1c). Hypoxia is also a risk factor for AD pathogenesis^33,34^, then we further assessed whether DIR is present in the blood of AD patients. Interestingly, AD patients had a high concentration of DIR in the blood (Fig. 1g; Supplementary Fig. S2a, b, d). Immunoblotting showed that DIR was present in blood samples from 85% of AD patients (n = 20) but only 16.7% of individuals from the general population (n = 18). The average density of the DIR band in the blood of AD patients (4992.38 ± 75.71, n = 20) was 5-fold higher than that in the blood of control subjects (931.58 ± 43.35, n = 18) and 3-fold higher than that in the blood of patients with GBM invading the hippocampus (1677.23 ± 40.10, n = 7), which was 2.5-fold higher than that in the blood of patients with GBM not invading the hippocampus (665.77 ± 40.29, n = 13). We also found that RNA level of DIR was significantly elevated in prefrontal tissue of AD patients by IRfinder^31^ (Supplementary Fig. S3a). On the other hand, we performed RT-PCR from hippocampus of control and AD patients, and found that *DIR* increased in AD patients (Supplementary Fig. S3b). Furthermore, DIR mainly colocalized with Aβ plaques in the hippocampus of AD patients (Fig. 1h). These results suggest that DIR could be translated and released in the hippocampus under pathological condition and then enter the blood of patients.

### Generation of homozygous DIR-KI mice

To test this hypothesis, we investigated the distribution and function of DIR in AD by generating DIR-KI mice using CRISPR/Cas9 technology. The DDIT4L protein is conserved in many species (Supplementary Fig. S4b). We compared the AA sequences of human and mouse DDIT4L and found that AAs 1-20 were identical and 2 AAs among AAs 21-30, corresponding to exon 2, were different, with the most marked difference being in the intronic region (Supplementary Fig. S4c). To induce the expression of human *DIR*, we inserted the sequence of the human *DIR* gene encoding the last 64 AAs after the sequence of the mouse *DDIT4L* gene encoding AAs 1-20 (Fig. 2a; Supplementary Fig. S4c). Genotyping and sequencing of newborn mice confirmed accurate sequence insertion (Fig. 2b). Of note, human DIR mRNA was amplified by designed primers (Fig. 2c). The body weight of the homozygous DIR-KI mice was not significantly different from that of the wild-type (WT) mice, suggesting that the mutant mice had normal growth and metabolism (Supplementary Fig. S5a). Moreover, the WT and homozygous DIR-KI mice showed no difference in distance travelled or velocity in the open field test, indicating that motor function was not altered (Supplementary Fig. S5b).

**Fig. 2:**
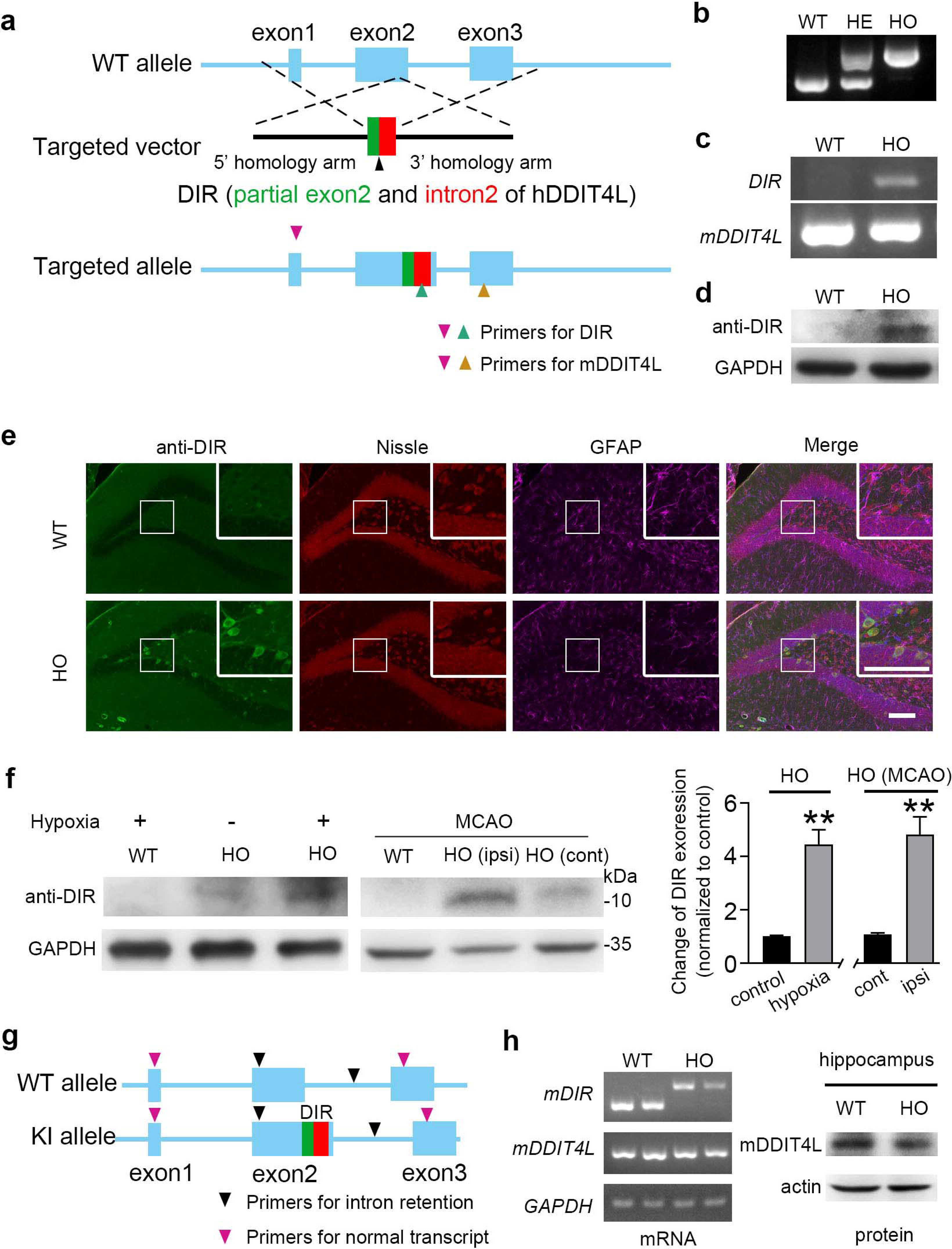
Generation of homozygous human DIR knock-in mice. a. The *DIR* knock-in (KI) mouse was constructed via CRISPR/Cas9 assays. To ensure the correct and intact expression of human *DIR*, the human *DIR* sequence encoding the last 64 AAs (the green block showing the 21-30 AAs of human DDIT4L, the red one for the 31-84 AAs of the DDIT4L intron translated region) was inserted behind the mouse *DDIT4L* genome that translated 1-20 AAs. b. Genotyping and sequencing of newborn mice confirmed the accurate sequence insertion of *DIR* (WT: wild-type, HE: heterozygote, HO: homozygote). c. The designed primers identified the human DIR mRNA in the HO mice. d. The DIR protein was expressed in the hippocampus of HO mice (n = 3). e. The DIR was expressed in the hippocampal dentate gyrus neurons of the HO mice. Scale bar = 100 mm. f. The immunoblotting showed the DIR expression in hippocampus was increased in the HO mice exposed to the hypoxia condition (8% O_2_, 8 h, n = 5), the DIR expression was apparently increased in the ipsilateral hippocampus in MCAO hypoxic-ischemic brain injury model of HO mice (n = 4). g and h. Specific primers confirmed the mouse intrinsic intron of DDIT4L (mDIR) in WT mice. The mDDIT4L expression have no change between WT and HO mice (n = 3).

Western blotting with an antibody targeting the IR portion of DIR showed the presence of one specific band (∼12 kDa) in the hippocampi of homozygous DIR-KI mice (Fig. 2d, Supplementary Fig. S6a). Immunostaining of brain slices from homozygous DIR-KI mice showed that DIR was expressed in many hippocampal dentate gyrus (DG) neurons (Fig. 2e; Supplementary Fig. S6b), some neurons in the CA1 and CA3 areas, and a few cortical and thalamic neurons. Therefore, the homozygous DIR-KI mice exhibited DIR protein expression in the hippocampus.

DIR expression in homozygous DIR-KI mice was assessed under hypoxic conditions. The expression of DIR was markedly enhanced in the hippocampi of homozygous DIR-KI mice exposed to low-oxygen conditions (8% O_2_) for 8 h (Fig. 2f). Unilateral middle cerebral artery occlusion (MCAO) was used to mimic hypoxic-ischemic brain injury, and we found that DIR expression was apparently increased in the ipsilateral hippocampi in homozygous DIR-KI mice subjected to MCAO but not in WT mice subjected to MCAO (Fig. 2f; Supplementary Fig. S6c). Thus, hypoxia could significantly enhance DIR expression *in vivo*.

To test whether the mouse intrinsic intron of DDIT4L could be retained, we designed specific primers to clone the fragment from mouse brain mRNA. Subsequent sequencing showed the existence of mouse DDIT4L intron retention (mDIR) mRNA in WT mice. The mDIR could not be transcribed in homozygous DIR-KI mice and not be translated in WT mice, while mDDIT4L expression was not changed in homozygous DIR-KI mice, as compared to that in WT mice (Fig. 2g, h, Supplementary Fig. S7a-d).

### AD-like pathological features in homozygous DIR-KI mice

To further investigate the potential roles of DIR in AD-like pathology, we performed immunostaining for Aβ and phosphorylated Tau (p-Tau), two classic AD-related molecules, in brain slices from 3-, 6-, and 9-month-old homozygous DIR-KI mice. The expression of Aβ and p-Tau was apparently upregulated and partly co-expressed with DIR in the hippocampi of homozygous DIR-KI mice (Fig. 3a; Supplementary Fig. S8a-d; Supplementary Fig. S9a-b; Supplementary Fig. S10a-b), while DDIT4L expression showed no change in homozygous DIR-KI and WT mice (Supplementary Fig. S10c). Taken together, the results indicate that DIR may initiate and promote the progression of AD pathology, particularly in the hippocampus.

**Fig. 3:**
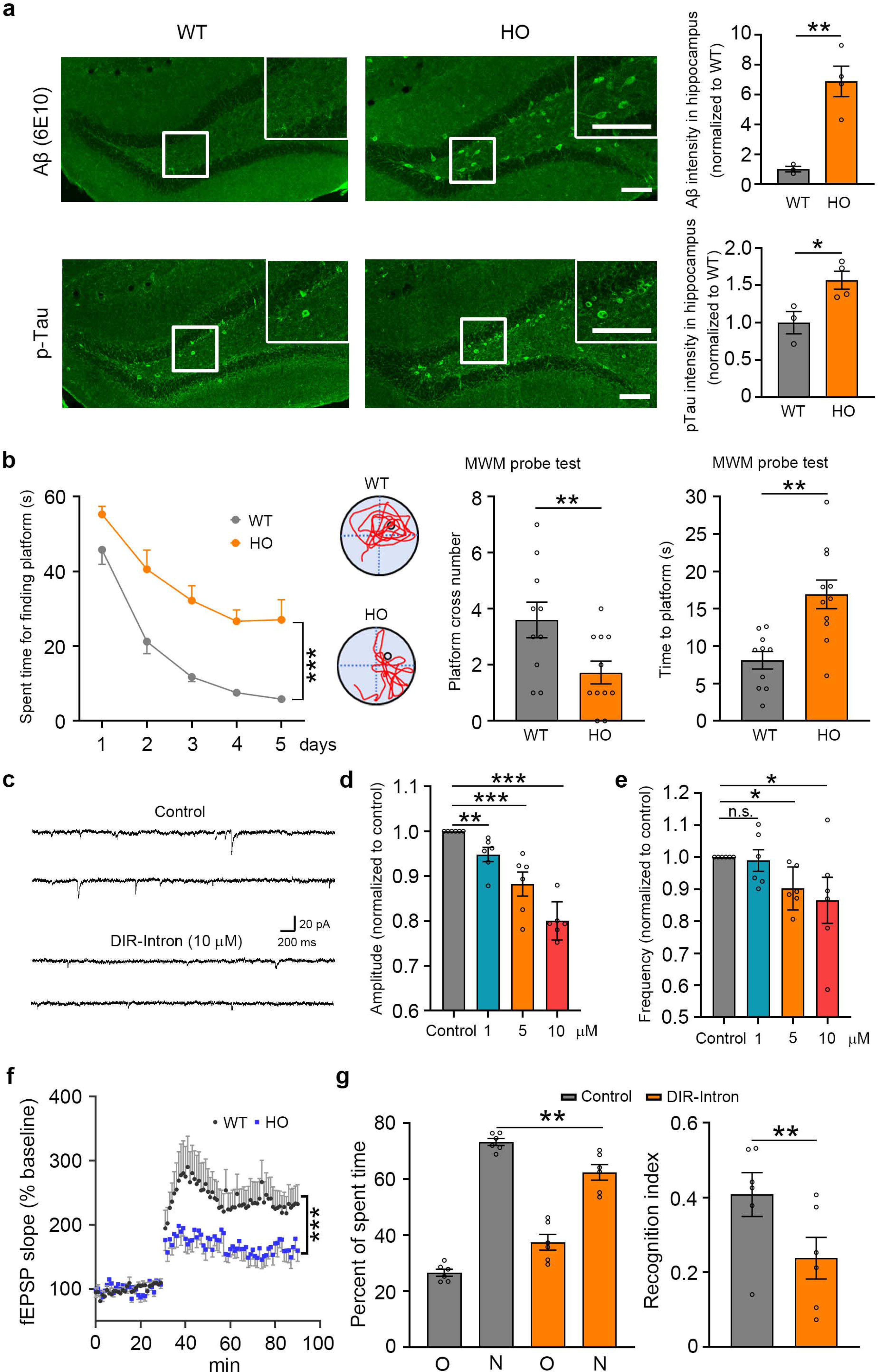
AD-like pathological features in homozygous DIR-KI mice. a. The expression of Aβ- or p-Tau was upregulated in the hippocampal dentate gyrus in the 6-month-old homozygous DIR-KI (HO) mice compared with wildtype (WT) mice. Scale bar = 100 μm. *, p < 0.05.**, p < 0.01. b. The Morris water maze test showed that the HO mice (n = 11) increased the escape latency following a 5-day training period compared with that of WT mice (n = 10). In the probe test to assess the spatial memory, WT mice had more times for crossing platform and spent less time to find the platform, compared with the HO mice. **, p < 0.01. c. The spontaneous excitatory postsynaptic current (sEPSC) of hippocampal neurons was inhibited by the DIR-Intron (10 mM, n = 6) in the brain slices prepared from WT mice. d. The amplitude of sEPSC in hippocampal neurons was suppressed by the DIR-Intron in a dose-dependent manner (n = 6). **, p < 0.01. ***, p < 0.001. e. The frequency of sEPSC in hippocampal neurons was dose-dependently reduced by the DIR-Intron (n = 6). *, p < 0.05. f. The long-term potentiation (LTP) of field EPSP recorded in the hippocampal CA1 region of the HO mice was apparently decreased, as compared with that of WT mice (n = 5 mice per group). ***, p < 0.001. g. The capacity of novel object recognition was reduced in the group treated with the DIR-Intron via the catheter implanted into the hippocampal CA1 region, delivering the DIR-Intron peptide locally through the subcutaneous minipumps (200 ng per mouse) (n = 6 mice per group). **, p < 0.01.

Next, we explored whether cognitive functions were impaired in homozygous DIR-KI mice. The results of the Y maze test showed that working memory was impaired in the homozygous DIR-KI mice (Supplementary Fig. S11a). Novel object recognition was attenuated in the homozygous DIR-KI mice (Supplementary Fig. S11b). In the Morris water maze test, most WT mice learned to use distal cues to navigate a direct path to the hidden platform following a 5-day training period. However, the homozygous DIR-KI mice showed an increased escape latency during the training process (Fig. 3b). In the probe test, which was used to assess spatial memory, the homozygous DIR-KI mice did not show a preference for the target quadrant, whereas WT mice showed a shorter latency to enter the target quadrant and crossed the platform more times (Fig. 3b).

To determine the functional domain of DIR, two polypeptides consisting of the first 30 AAs (DIR-Exon) or the last 54 AAs (DIR-Intron) of DIR were synthesized. Hippocampal slices from WT mice were incubated with either DIR-Exon or DIR-Intron (1, 5 and 10 μM). Notably, only DIR-Intron (1, 5 μM and 10 μM) dose-dependently reduced the spontaneous excitatory postsynaptic current (sEPSC) amplitude and frequency, whereas neither the amplitude nor the frequency of sEPSCs was altered after DIR-Exon incubation (Fig. 3c-e; Supplementary Fig. S11c, d). Moreover, long-term potentiation (LTP) of field excitatory postsynaptic potentials (EPSPs) recorded in the hippocampal CA1 region was apparently decreased in brain slices from homozygous DIR-KI mice compared with those from WT mice, indicating that DIR suppresses the major excitatory synaptic pathway in the hippocampus (Fig. 3f). Furthermore, we found that local delivery of the peptide DIR-Intron (200 ng per mouse) via osmotic pumps connected to catheters inserted into the CA1 region of the hippocampus decreased novel object recognition (Fig. 3g). Overall, the homozygous DIR-KI mice displayed progressive Aβ deposition as well as increased Tau phosphorylation coinciding with the decline in cognition and memory because of the inhibitory effect of DIR on excitatory synaptic transmission in hippocampal neural circuits.

To test whether the mouse intrinsic intron of DDIT4L contributes to learning and memory, we designed specific siRNA for mDIR (Supplementary Fig. S12a, b). The siRNA did not affect the behavior performance in the Y maze test or novel object recognition in WT or APPPS1 mice (Supplementary Fig. S12c, d), indicating that mDIR is not involved in learning and memory processes.

### DIR induces Aβ deposition via gelsolin

In AD patient samples, DIR mainly colocalized with Aβ in the hippocampus (Fig. 4a, Supplementary Fig. S13a). The interaction between DIR and Aβ was observed in transfected HEK293 cells, but DDIT4L (including DIR-Exon) could not interact with Aβ (Fig. 4b; Supplementary Fig. S14a). These results indicate that DIR-Intron could be the main region of DIR that interacts with Aβ. Then, we investigated whether DIR-Intron and Aβ directly interact. Proximity ligation assays (PLAs) showed that DIR did not directly interact with Aβ (Fig. 4c). Using synthetic DIR-Intron and Aβ proteins, we found that DIR-Intron could not directly bind to Aβ (Supplementary Fig. S14b), suggesting that DIR has another binding target. Then DIR was transfected into human brain cell lines, and the cell lysates were immunoprecipitated with an anti-DIR antibody, one immunoreactive band with a molecular weight of ∼85 kDa was coimmunoprecipitated (Fig. 4d). The band was extracted and further analysed by mass spectrometry. This analysis identified 12-15 peptides that matched the human gelsolin protein (∼85 kDa), covering 35% of the gelsolin sequence (Fig. 4e).

**Fig. 4:**
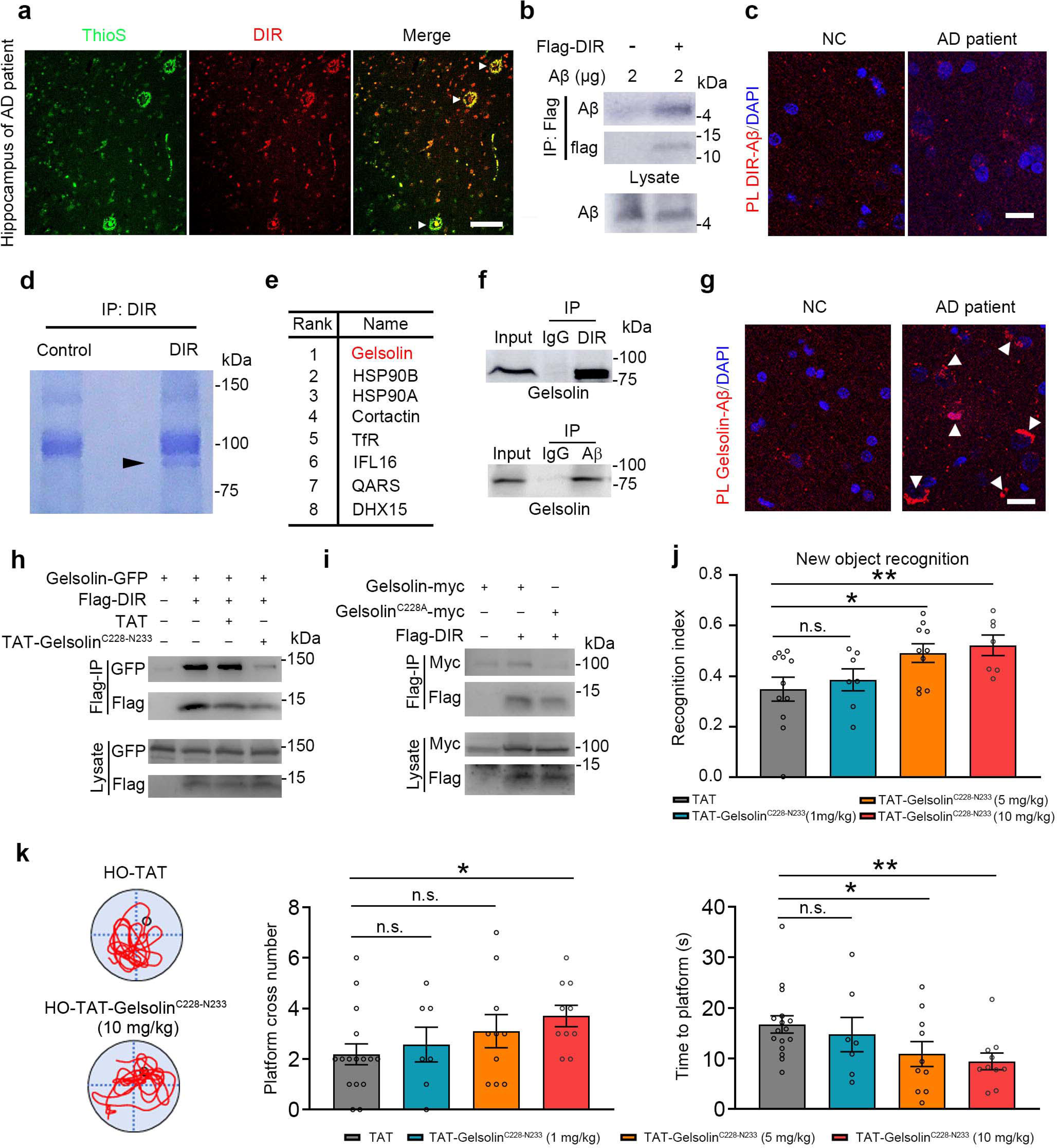
DIR induces Aβ deposition via gelsolin. a. DIR was present in the thioflavine S (ThioS)-positive plaque (arrows) in the hippocampus of AD patients (n = 3). Scale bar = 50 mm. b. In the lysate of HEK293 cells transfected with the plasmid expressing Flag-DIR with an addition of synthetic Ab42, Ab was found in the proteins precipitated with Flag antibodies (n = 3). c. The PLA assay showed that DIR could not bind to Aβ in AD patients’ hippocampus. Scale bar = 50 mm. d. Coomassie blue staining showed a band of molecular weight ∼85 kDa in the DIR-antibody-precipitated proteins from U87MG cells transfected with the Flag-DIR plasmid (n = 3). e. The mass spectrometry identified top 8 molecules from the ∼85 kDa band. Gelsolin was the most abundant molecule in the list. f. In the lysate of hippocampus from the DIR-KI mice, gelsolin was found in the proteins precipitated with the DIR and Ab antibodies (n = 3). g. The PLA assay showed that gelsolin bound to Ab (arrows) in AD patients’ hippocampus. Scale bar = 50 mm. h. In the lysate of HEK293 cells co-transfected with plasmids expressing Flag-DIR and Gelsolin-GFP, Gelsolin-GFP was found in the proteins precipitated with Flag antibodies. The TAT-Gelsolin^C228-N233^ apparently reduced the DIR/Gelsolin interaction (n = 3). i. In the lysate of HEK293 cells co-transfected with plasmids expressing Flag-DIR and Gelsolin-myc or Gelsolin^C228A^-myc, Gelsolin-myc, not Gelsolin^C228A^-myc, were found in the proteins precipitated with Flag antibodies (n = 3). j. The capacity of novel object recognition was reversed by the TAT-Gelsolin^C228-N233^ treatment in a dose-dependent manner in the HO mice (1 mg/kg, n = 7; 5mg/kg, n =10; 10 mg/kg, n = 7), as compared with the mice treated with TAT (n = 11). *, p < 0.05. **, p < 0.01. k. Representative trial traces of individual mice on probe test day. In the probe test to assess spatial memory, the mice treated with the TAT-Gelsolin^C228-N233^increased time of crossing platform and spent less time to find platform in a dose-dependent manner (1 mg/kg, n = 7; 5mg/kg, n =10; 10 mg/kg, n = 10), compared with the mice treated with TAT (n = 16). *, p < 0.05. **, p < 0.01.

Gelsolin has been shown to interact with Aβ^35^, forming a complex that is transported to the circulatory system^36-38^. Our study showed that DIR interacted with gelsolin in DIR-KI mice, HEK293 cells and human tissues (Fig. 4f, g; Supplementary Fig. S14c). In the lysates of HEK293 cells that were transfected with DIR and endogenously expressed gelsolin, the addition of Aβ42 enhanced the interaction of DIR with gelsolin in a dose-dependent manner, indicating that Aβ contributes to the interaction between DIR and gelsolin (Supplementary Fig. S14d). To determine the interaction domain of gelsolin, we checked a possible docking site for DIR in gelsolin using a combination of homology modelling and in silico docking^39-41^. In this model, we screened C228-N233 motifs in gelsolin (Gelsolin^C228-N233^) as possible docking sites for DIR (Supplementary Fig. S14e). The interaction between DIR and gelsolin could be disturbed by the synthesized TAT-gelsolin^C228-N233^ peptide (Fig. 4h). Moreover, C228 in gelsolin is the key point for the interaction between DIR and gelsolin (Fig. 4i). Interestingly, the DIR-KI mice, given TAT-gelsolin^C228-N233^, showed obvious cognitive improvement in new object recognition and Morris water maze in a dose-dependent manner (Fig. 4j, k), indicating that Aβ contributed cognitive impairment in DIR-KI mice model.

### DIR contributes Aβ plaque formation

Moreover, the synthetic DIR-Intron and Aβ42 proteins were added to purified gelsolin or HEK293 cell lysates, and DIR-Intron apparently enhanced the formation of insoluble Aβ42, which is a major constituent of amyloid plaques (Fig. 5a; Supplementary Fig. S16a). Therefore, the intron region of DIR is involved in Aβ deposition. It is known that thioflavine S is a specific marker of Aβ plaques in the brains of AD patients^42^. However, thioflavine S did not label the amyloid plaques in the hippocampus of mice because the murine Aβ sequence cannot be detected by thioflavine^43^. Interestingly, the slices of hippocampal sections from DIR-KI mice that were incubated with human Aβ42 or Aβ40 peptides for 6 h could be stained by thioflavine S (Fig. 5b; Supplementary Fig. S15a), which was consistent with the thioflavine S staining of human Aβ plaques. Importantly, triple immunostaining showed that DIR was colocalized with gelsolin and Aβ to a greater degree in dense-core plaques than in diffuse plaques in the hippocampus and cortex regions of AD patient, while there was no obvious DIR or amyloid plaque staining in the sections from control subjects without AD (Fig. 5c; Supplementary Fig. S15b). Moreover, 3D imaging, thioflavine T and electron microscope showed DIR bound gelsolin inducing Aβ deposition (Supplementary Fig. S15C and Fig. S16b and Fig. S16c). Taken together, these results show that an increase in DIR expression could lead to Aβ deposition and subsequent plaque formation by binding to gelsolin under pathological conditions. Therefore, DIR may be an inducer of Aβ deposition and plaque formation.

**Fig. 5:**
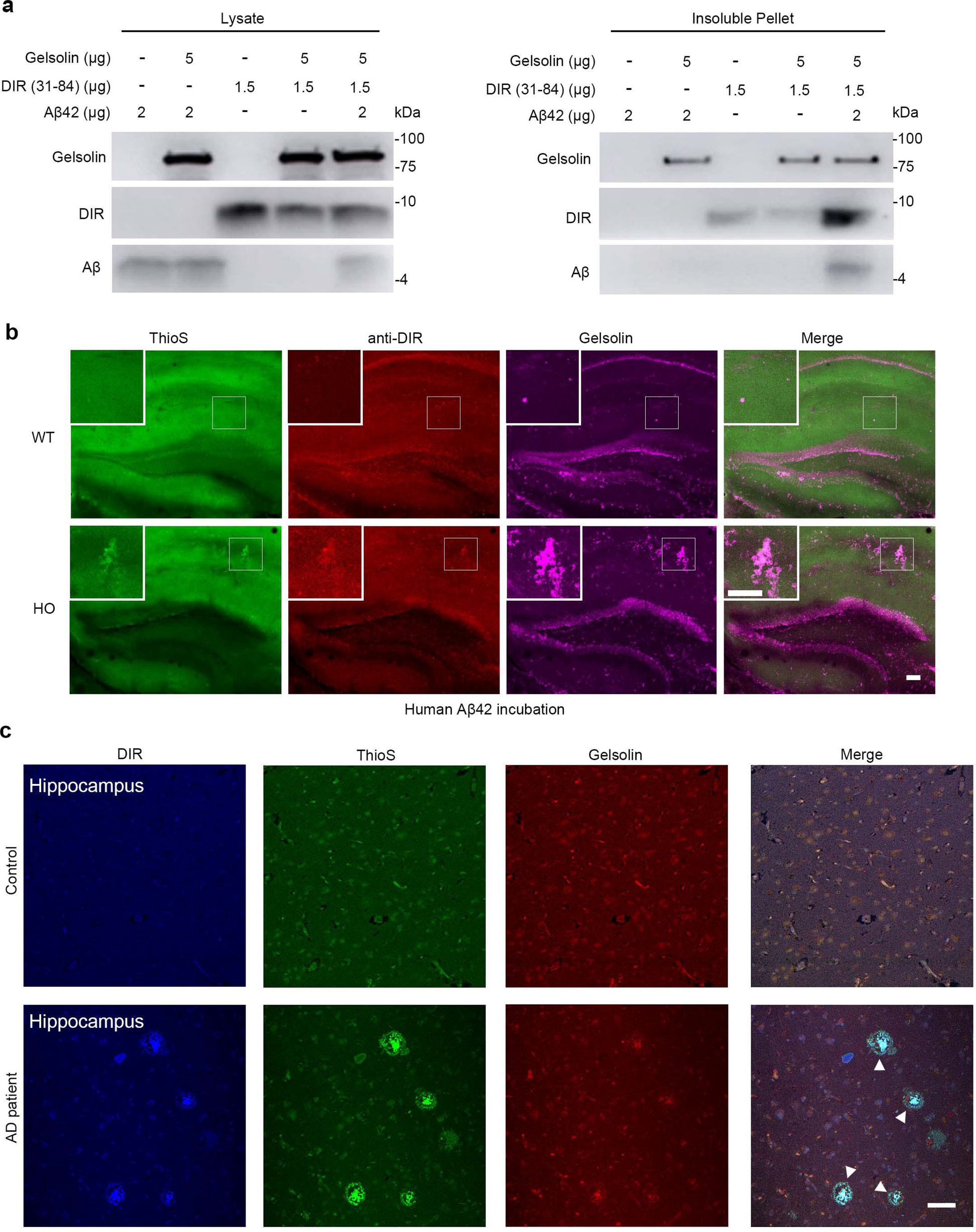
DIR mediates Aβ plaque formation. a. In the mixture of gelsolin-his with synthetic Aβ42 or DIR (31-84), DIR (31-84) induced Aβ42 insolubilization via gelsolin (n = 3). b. In the hippocampus slices of the homogenous DIR-KI (HO) mice (n = 3), but not that of wildtype (WT) mice (n = 3), incubated with the synthetic human Aβ42 (100 μM) for 6 h, thioflavine S (ThioS)-positive plaque (arrow) appeared and co-existed with DIR and gelsolin. Scale bar = 100 μm. c. In AD patients (n = 3), but not control individuals (n = 3), ThioS-positive dense-core plaque co-localized with DIR and gelsolin (arrows) in the hippocampal DG area. Scale bar = 50 μm. Quantitative analysis showed that DIR, gelsolin and ThioS-positive in 86.3% dense-core plaque in cerebral cortex, in 92.8% dense-core plaque in hippocampus.

### DIR binding to the GluA1 subunit of the AMPA receptor induces cognitive impairment and synaptic deficiency

Next, we further explored whether there was a new target for DIR in AD. The hippocampal lysates of homozygous DIR-KI mice were immunoprecipitated with an anti-DIR antibody. One immunoreactive band with a molecular weight of ∼110 kDa was coimmunoprecipitated (Fig. 6a). The band was extracted and further analyzed by mass spectrometry. This analysis identified 13 peptides that matched the human GluA1 protein, covering 14% of the GluA1 sequence (Fig. 6b). Our study showed that DIR interacted with GluA1 in DIR-KI mice (Fig. 6c, d). To determine the precise binding region of GluA1, we investigated the key motifs that were involved in the DIR modulation of AMPA receptor activation. We first checked a possible docking site for DIR in GluA1 using a combination of homology modelling and in silico docking^39-41^. In this model, we screened R198-E205 motifs in GluA1 (GluA1^R198-E205^) as possible docking sites for DIR (Fig. 6e). The interaction between DIR and GluA1 could be disturbed by the synthesized TAT-GluA1^R198-E205^ peptide (Fig. 6f). Moreover, C204 in GluA1 is the key point for the interaction between DIR and GluA1 (Fig. 6g).

**Fig. 6:**
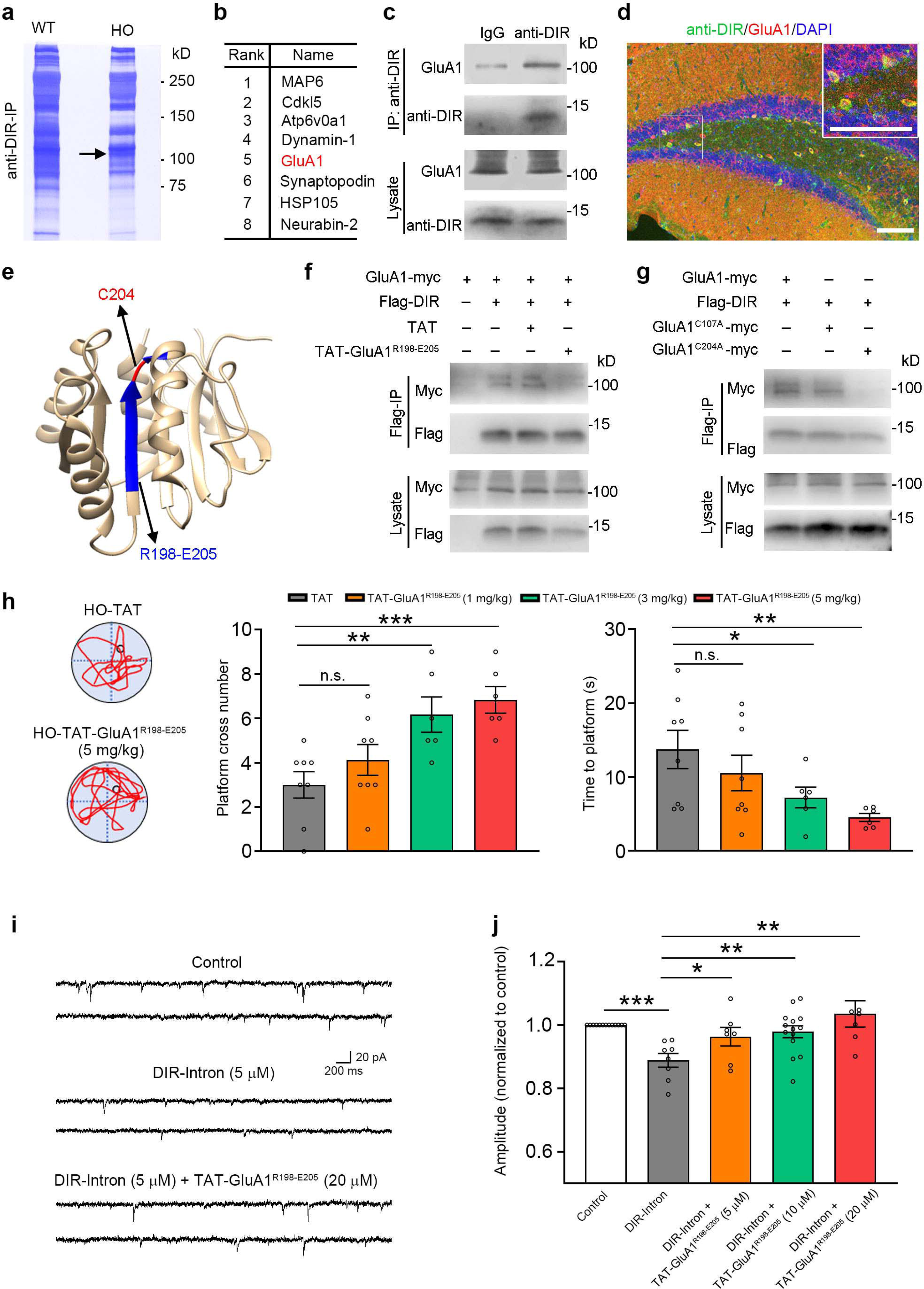
DIR/GluA1 interaction induces cognitive impairment and synaptic deficiency. a. Coomassie blue staining showed a band of molecular weight ∼110 kDa in the DIR-antibody-precipitated proteins from the hippocampus of homogenous DIR-KI (HO) mice, but not wildtype (WT) (n = 3). b. The mass spectrometry identified top 8 molecules from the ∼110 kDa band. GluA1 was in the list. c. In the lysate of hippocampus from the HO mice, GluA1 was found in the proteins precipitated with the DIR antibody (n = 3). d. The immunostaining showed that DIR was colocalized with GluA1 in the HO mice. Scale bar = 50 mm. e. The R198-E205 motif in GluA1(GluA1^R198-E205^) could be the docking site for DIR. C204 in GluA1 could be the key point for the interaction between DIR and GluA1. f. In the lysate of HEK293 cells co-transfected with plasmids expressing Flag-DIR and GluA1-myc, GluA1-myc was found in the proteins precipitated with Flag antibodies. The TAT-GluA1^R198-E205^ apparently reduced the DIR/GluA1 interaction (n = 3). g. In the lysate of HEK293 cells co-transfected with plasmids expressing Flag-DIR and GluA1-myc or GluA1^C107A^-myc or GluA1^C204A^-myc, GluA1-myc and GluA1^C107A^-myc, not GluA1^C204A^-myc, were found in the proteins precipitated with Flag antibodies (n = 3). h. Representative trial traces of individual mice on probe test day. In the probe test to assess spatial memory, the HO mice treated with TAT (n = 8) did not show a preference for the target quadrant, whereas the mice treated with the TAT-GluA1^R198-E205^ had more times for crossing platform and spent less time to find platform in a dose-dependent manner (1 mg/kg, n = 8; 3 mg/kg, n = 6; 5 mg/kg, n = 6). *, p < 0.05. **, p < 0.01. ***, p < 0.001. i. The spontaneous excitatory postsynaptic current (sEPSC) of hippocampal neurons was inhibited by the DIR-Intron (5 mM, n = 8). TAT-GluA1^R198-E205^ (20 mM, n = 6) reversed this inhibition in the brain slices prepared from WT mice. j. The amplitude of sEPSC in hippocampal neurons was suppressed by the DIR-Intron. TAT-GluA1^R198-E205^ could reverse this suppression in a dose-dependent manner (5 mM, n = 7; 10 mM, n = 14; 20 mM, n = 6). *, p < 0.05. **, p < 0.01. ***, p < 0.001.

To examine whether GluA1^R198-E205^ could alleviate AD-like pathogenesis by binding DIR, we intraperitoneally injected TAT-GluA1^R198-E205^ (1, 3, 5 mg/kg) or control peptide into homozygous DIR-KI mice for 7 days. TAT-GluA1^R198-E205^ peptide treatment decreased the escape latency during the training phase of the Morris water maze test. In the probe test the mice treated with TAT-GluA1^R198-E205^ crossed the platform more times and spent less time to find platform (Fig. 6h). The inhibitory effect of DIR-Intron (5 μM) on sEPSCs could be reversed by TAT-GluA1^R198-E205^ (5, 10, 20 μM) (Fig. 6i, j). The decrease in hippocampal LTP in the homozygous DIR-KI mice could be rescued by intraperitoneal injection of TAT-GluA1^R198-E205^ (3 mg/kg, 7 days) (Supplementary Fig. S17a). However, TAT-GluA1^R198-E205^ treatment did not change the expression of Aβ and p-Tau in the hippocampi of homozygous DIR-KI mice (Supplementary Fig. S17b).

### Alleviating AD-like pathological features with a DIR mAb

To examine whether a mAb against DIR could alleviate AD-like pathogenesis, we intraperitoneally injected the DIR mAb (0.5 mg/day) or IgG into homozygous DIR-KI mice for 5 days. Both working memory and novel object recognition were improved by mAb treatment (Supplementary Fig. S18a, b). The mAb treatment decreased the escape latency during the training phase of the Morris water maze test. In the probe test, the mice treated with the mAb had a shorter latency to enter the target quadrant and crossed the platform more times (Fig. 7a). Moreover, mAb could penetrate the blood-brain barrier and reduced the expression of Aβ and p-Tau in the hippocampi of homozygous DIR-KI mice (Fig. 7b, c; Supplementary Fig. S18c). The decrease in hippocampal LTP in the homozygous DIR-KI mice could be rescued by intraperitoneal injection of the mAb (0.5 mg/day, 5 days) (Fig. 7d). Hippocampal slices from DIR-KI mice intraperitoneal injected with the DIR mAb showed increased sEPSC amplitude, compared with those from DIR-KI mice intraperitoneal injected with IgG (Fig. 7e-g). Collectively, these results indicate that inhibiting DIR activity with a DIR mAb not only decreased the levels of Aβ and p-Tau but also reversed cognition and memory deficits in homozygous DIR-KI mice, indicating the important role of DIR in AD-like pathogenesis.

**Fig. 7:**
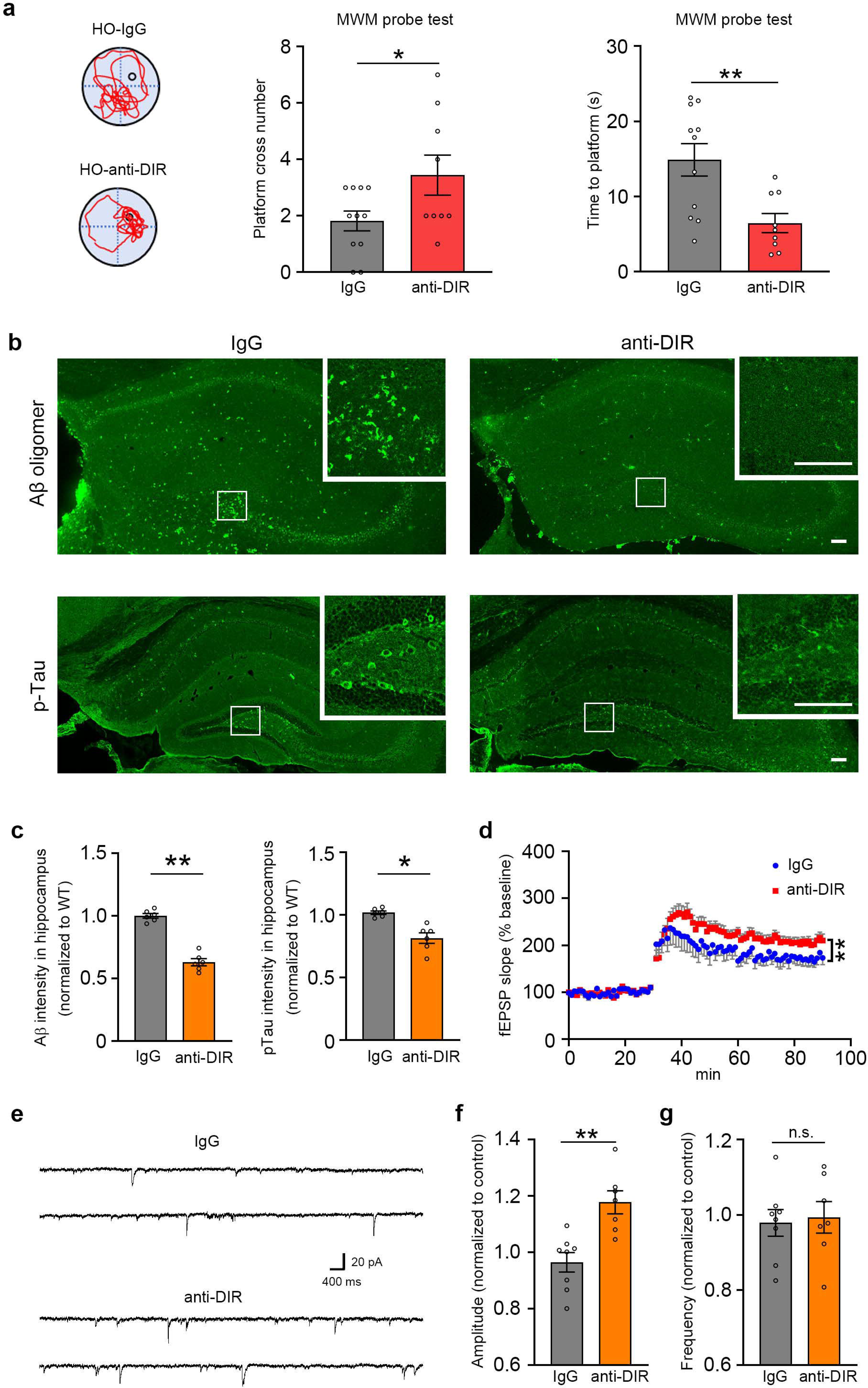
Reversing AD-like pathological features by DIR mAb. a. Representative trial traces of individual mice on probe test day. In the probe test to assess spatial memory, the mice treated with the DIR mAb (n = 9) had more times for crossing platform and spent less time to find platform, compared with the mice treated with IgG (n = 11). *, p < 0.05. **, p < 0.01. b. The presentation of Aβ- and p-Tau expression in the hippocampal dentate gyrus in the HO mice treated with the DIR mAb. Scale bar = 100 μm. c. The expression of Aβ- and p-Tau was apparently decreased in the hippocampal dentate gyrus in the HO mice treated with the DIR mAb. *, p < 0.05. **, p < 0.01. d. The LTP in theHO mice was apparently increased after the DIR mAb treatment, as compared with IgG treatment (n = 5 mice per group). **, p < 0.01. e. The spontaneous excitatory postsynaptic current (sEPSC) of hippocampal neurons was increased after the DIR mAb treatment (n = 7), as compared with IgG treatment (n = 8). f. The amplitude of sEPSC in hippocampal neurons was apparently increased after the DIR mAb treatment (n = 7), as compared with IgG treatment (n = 8). **, p < 0.01. g. The frequency of sEPSC in hippocampal neurons had no change after the DIR mAb treatment (n = 7), as compared with IgG treatment (n = 8).

## Discussion

Briefly, we identified the presence of a new pathogenic molecule, DIR, which results from IR due to aberrant splicing of *DDIT4L*, in AD patients. DIR*-*carrying mice exhibited accumulation of Aβ and p-Tau in the hippocampus as well as decreased cognition and memory, which correlated with the suppressive effect of DIR on hippocampal LTP. These findings demonstrate the pathogenic role of hypoxia-induced DIR in the contribution to dementia and show that homogenous DIR-KI mice can be used as a model for dementia pathology. Furthermore, inhibition of DIR function in the homogenous DIR-KI mice alleviated cognitive decline, suggesting the potential of DIR as a target for the development of dementia therapies.

Previous studies indicated that gelsolin would prevent or delay the progression of AD by sequestering Aβ and inhibiting Aβ plaque formation. Our study showed gelsolin was an important linker between DIR and Aβ deposition. Disturbed interaction of gelsolin and DIR via TAT-gelsolin^C228-N233^, which led to Aβ clearance from brain. This study also presented DIR induced cognitive impairment, such as novel object recognition and spatial learning and memory, had been reversed by TAT-gelsolin^C228-N233^. Our results indicated soluble Aβ peptides were present in the peripheral zone while DIR was present in central zone of dense-core plaques in AD patients. This indicated that neural transmission would be inhibited by soluble Aβ after dense-core plaques formation, which may account for behavior changes.

On the other hand, AMPA receptors in the hippocampus are key sites for synaptic LTP, learning and memory^44-46^. In this study, we identified a new protein, DIR, that bound the GluA1 subunit of AMPA receptors, inhibited synaptic transmission and induced cognitive impairment. The DIR protein sequence is unique in humans, not in mice and rats. Only under pathological conditions, such as hypoxia, could its expression be induced. Moreover, no obvious mDIR protein expression in mice under hypoxia may also reveal the specificity of the intron sequence of DIR, which needs further investigation. These conditions enhance the difficulty of identifying the role of DIR in cognitive impairment, especially in AD. The mono-antibody of DIR could reverse DIR-induced dementia pathological features. More interestingly, the synthetic binding domain of GluA1 for DIR could disturb the interaction of DIR and GluA1, and reverse cognitive impairment. This indicates that the DIR-GluA1 pathway is an important factor for cognitive impairment.

The splice of IR may be contributed to pathogenesis of neurodegenerative disease^30,47^. In this study, DIR had been identified as an IR form of DDIT4L. As mapping the intron sequence of *DDIT4L* RNA, homologous degree reveals their evolutionary relationship in different species. Human and chimpanzee have similar sequence except for two points in the identified retention sequence, while the sequences in human and mouse have the least similarity in all compared species (supplementary table 1). Therefore, the mouse is the best model to check DIR function. On the other hand, IR appears to be mostly directly related to the splicing pathway involving the formation of spliceosome for the removal introns^48,49^. While in human, spliceosome function has stronger than other species for human proteome complexity^50^. In our study, DIR in human is not expressed in normal condition while DIR in DIR-KI mice has been existed in normal condition, this difference may be determined by spliceosome function in different species.

Cerebrovascular sclerosis and other pathological conditions that lead to hypoxia often occur in elderly individuals^19,21,34^. The subsequent hypoxia in local microcerebrovascular circulation may result in abnormal intron retention^25,29^, including the translation of the DIR protein. Then, DIR bound to gelsolin and induced Aβ deposition to form dense-core plaques in the brain. Moreover, DIR interacted with GluA1, contributing to synaptic deficiency and cognitive impairment. Therefore, DIR would be a novel potential therapeutic target for dementia. Furthermore, TAT-gelsolin^C228-N233^, TAT-GluA1^R198-E205^ and DIR mAb inhibit DIR-induced cognitive decline, suggesting that TAT-gelsolin^C228-N233^, TAT-GluA1^R198-E205^ and DIR mAb could be therapies for cognitive impairment in dementia.

## Materials and methods

### STAR METHODS

### Human samples

The human brain samples collection was conducted with the approval of the Ethics Committee of Huashan Hospital of Fudan University. Informed consent was obtained from all subjects involved in this study. In detail, the cDNA of patients (L1002 and G2913) was used for the detection of hDDIT4L mRNA (forward primer: 5’-TGCTGGACTGTGGCTATCAC-3’ and reverse primer: 5’-ACAAGGACCTTTGAGCAACCA-3’). The tissue lysates of patients (L4023, G2913 and G1359) were used for the examination of hDDIT4L and DIR proteins. The MRI brain images of GBM patients (G8098 and G8409) were used for the definition of tumor infiltration. The plasma of GBM, LGG patients and AD patients were used to examine the DIR. The information of enrolled patient was supplied in supplementary table 1.

### Homozygous DIR-KI mice

All animal operations were under the guidelines of the National Center for Protein Science Shanghai. Mice were accommodated in the specific pathogen-free rooms and supplied with enough foods and water. The DIR-KI mice were generated by Shanghai Model Organisms Center, Inc. Briefly, a cDNA encoding human *DIR* was inserted into mouse *DDIT4L* gene locus (Fig. 2a and Supplementary Fig. 2c) through homologous recombination by using CRISPR/Cas9 technology. All mice were maintained on a C57BL/6J background. The neonatal mice were identified by the PCR analysis, using the following genotyping primers.

Forward primer: 5’-GAGGAGCCTGTGCACTTCTT-3’;

Reverse primer: 5’-CACACCCAGCCTGTACACTT-3’.

The adult mice were anaesthetized and sacrificed. Then the brain tissues were isolated and homogenized. Next, total mRNAs were extracted using Trizol regents and 1 μg mRNAs were transcribed reversely to cDNA.

The following primers were used to detect the expression of DIR mRNA.

Forward primer: 5’-TGCCTCGGTTTACCCTTC-3’;

Reverse primer: 5’-AAGATGTTAGAAAATTTGGAAAGG-3’.

The following primers were used to detect the expression of mDIR mRNA.

Forward primer: 5’-ACGGGCAGTTTGAGCAGTAA-3’;

Reverse primer: 5’-CACAAGTAAGCGCATGTCTTT-3’.

The following primers were used to detect the expression of mDDIT4L mRNA.

Forward primer: 5’-TGCCTCGGTTTACCCTTC-3’;

Reverse primer: 5’-CCAGCTTTTTACATACATTTTCA-3’.

The following primers were used to detect the expression of GAPDH mRNA.

Forward primer: 5’-TATGTCGTGGAGTCTACTGGTGTCTTCACC-3’;

Reverse primer: 5’-GTTGTCATATTTCTCGTGGTTCACACCC-3’.

### Antibodies

The DDIT4L and DIR antibodies used in our research were produced by GL Biochem (Shanghai) Ltd. The polyclonal antibodies used for Western blotting were generated from the rabbits, while the useful monoclonal antibodies for immunostaining and intraperitoneal injection were generated from the hybridoma.

To test the specificity of antibodies, the antibodies and the antibodies pre-absorbed with the antigens (10^-5^ or 10^-6^ M) for 12 h at 4□ were used to incubate the nitrocellulose membranes containing the same lysate samples, respectively. The nitrocellulose membranes were then incubated with HRP ligated second antibodies and imaged. Moreover, the antibody specificity also tested with the immunostaining of brain sections of homozygous DIR-KI mice.

### Immunoprecipitation

The HEK293 cells and brain tissue of DIR-KI mice were lysed in the ice-cold buffer (25 mM Tris, 150 mM NaCl, 1 mM EDTA, 1% NP40, 5% glycerol and protease inhibitors, PH = 7.5). The suspended lysate was immunoprecipitated with the antibodies overnight at 4□ and then added Protein G-Agarose to bind the antibodies for 4 h at 4□. The sepharose was then resuspended in the RIPA buffer without SDS, washed at least 3 times, and incubated in the SDS buffer for 20 min at 60□. Then the immunoblotting was processed. The immunoblot band in DIR-KI mice was analyzed with the mass spectrometry.

### Immunoblotting

Adult mice were anaesthetized and perfused with the PBS. Then brain tissues were extracted and homogenized on ice in the RIPA buffer (100 mM Tris, 150 mM NaCl, 1% Triton X-100, 10% glycerol, 0.05% BSA, 0.1% SDS and protease inhibitors, PH = 7.5). After centrifuged, supernatants of brain lysates were collected for further experiments. The brain tissues of human were lysed with the same tissue RIPA buffer. The plasma of human was first diluted using the PBS at a ratio 1:10, about 5 μl samples of each diluted plasma were loaded to gels. All samples were fractionated by the sodium dodecyl sulfate (SDS)-gel electrophoresis, followed by being transferred onto the nitrocellulose filter membranes, and blocked with 5% nonfat milk in the TBS buffer containing Tween-20 (TBST). The membranes were incubated in the primary antibodies overnight at 4°C. Next day, membranes were washed with the TBST for three times and incubated with the HRP ligated second antibodies. After additional washing, the membranes were prepared to be imaged by adding the ECL buffer. Protein bands were imaged by using an imaging system (GE). The primary antibodies used were listed here: Flag (Sigma, SAB4200071), Myc (Cell Signaling Technology, 2276S), Actin (Chemicon, MAB1501), Tubulin (Sigma, T5168), GAPDH (Proteintech, 10494), Gelsolin (Abcam, ab109014), GluA1 (Synaptic Systems, 182011), Aβ (Cell Signaling Technology, 8243S), and GFP (Invitrogen, A11122).

### Immunohistochemistry

Briefly, mice were anaesthetized and perfused with Lana’s fix buffer or 4% PFA. Then brain tissues were collected and fixed in new fix buffer at 4°C for 1 h. After washing in PBS for three times, tissues were placed in 30% sucrose/PBS overnight at 4°C, then mounted in the OCT compound. Frozen section was performed to obtain brain section (40 µm in thickness). For immunostaining, tissue sections were blocked with PBST (0.1% Triton X-100/2.5% normal donkey serum/PBS) for 30 min at room temperature, followed by the incubation with primary antibodies overnight at 4°C. Then tissue sections were washed with PBS for three times and incubated with Alexa fluorescence-conjugated secondary antibodies (Invitrogen) for 60 min at 37°C, then washed with PBS three times and mounted with medium containing DAPI. The images were acquired using a Leica SP8 confocal microscope. The primary antibodies used were listed here: GFAP (Sigma, G9269), Aβ (BioLegend, 803001), Aβ oligomer (Invitrogen, AHB0052), p-Tau (Cell Signaling Technology, 12885), GluA1 (SYSY, 182011).

Paraffin section of human brain tissues were provided by: Human Brain Bank, Chinese Academy of Medical Sciences & Peking Union Medical College, Beijing, China. Immunofluorescence staining was used to study the distribution of DIR and Aβ. In a single tissue section, primary antibodies raised in different species (mouse, rabbit or sheep) were used. The secondary antibodies (anti-mouse, anti-rabbit or anti-sheep) were coupled to different fluorochromes (Alexa Fluor 488, Cy3 or Cy5). After staining, cover the fluorescently labeled sections with VECTASHIELD mounting medium (Vector Labs). The primary antibodies and dye used were listed here: Aβ (Cell Signaling Technology, 8243S), Thioflavin S (ChemCruz, sc-391005), Gelsolin (Abcam, ab109014).

### Magnetic resonance imaging (MRI)

The MRI brain images were acquired with a 3.0T system of MRI. A high-resolution anatomical volume image was obtained using the following parameters: TR (repetition time)/TE (echo time) = 2300/2.27 ms, inversion time = 900 ms, slice thickness = 1.0 mm, FoV (field of view) read = 250 mm, FoV phase = 100%, PAT (parallel acquisition technique) = 3, TA (acquisition time) = 3:54 min.

### Analysis of differential gene expression

The Cancer Genome Atlas (TCGA) diffuse glioma database, including the RNA-seq, simple somatic mutation and patient survival, were downloaded from the TCGA Research Network by R Studio Version 1.2.5033 with packages TCGA biolinks^51^. The expression of DDIT4L and other genes in the TCGA diffuse glioma database were processed using the publicly available tools, including the biostatistics program R Studio with packages gplots, DT, dplyr and Summarized Experiment.

### RNA-seq Data and processing

The Alzheimer’s disease human brain transcriptomes were obtained from the National Center for Biotechnology Information (NCBI). The accession number is GSE153873 and GSE159699^52^. The tissue type all is postmortal lateral temporal lobe. The average sequencing depth of the subjects was 25 to 30 million reads. The data were aligned to the human reference genome (version 110 of GRCm38) using STAR^53^.

### Quantification of intron retention using IRFinder

For using IRFinder^54,55^, we first built a reference using ‘IRFinder BuildRef’ command based on the Release 110 of Ensembl human genome GRCh38 gene annotations, including RNA.SpikeIn.ERCC.fasta.gz and Human_hg38_nonPolyA_ROI.bed files obtained from IRFinder repository (https://github.com/RitchieLabIGH/IRFinder). After downloading the transcriptome FASTQ files as described in the previous subsection, we quantified the intron retention via ‘IRFinder FastQ’ command. We used the values of ‘SpliceExact’ column as a measure for intron events.

### NPCs and neuronal differentiation

Human embryonic stem (ES) cells were used to be differentiated into neural progenitor cells (NPCs) and then differentiated into neurons. The NPCs were maintained in Dulbecco’s modified eagle’s medium: F12 (DMEM: F12, Thermo Fisher) containing 1×B27 (Sigma-Aldrich), 1×N2 (Gibco), and basic fibroblast growth factor (20 ng/mL, bFGF, Sigma-Aldrich) at 37°C with 5% CO_2_. The NPCs were differentiated into neurons in the absence of bFGF for at least 3 weeks and treated with 10 μM ROCK inhibitor (Y-27632; Calbiochem) during the first 24 h.

### Plasmid construction and expression

Primers, inducing the extra homology arms paired with the pCMV-flag vector constructed based on the pCMV-myc vector, were designed to clone the full length of DIR CDS sequences.

Forward: 5’-CCATGGAGGCCCGAATTCGGATGGTTGCAACTGGC-3’;

Reverse: 5’-ACTCATCAATGTATCTTATCTTATGGAGAGAAGATGTTAGAAAA- 3’.

Primers, inducing the extra homology arms paired with the pcDNA3.1-myc-his vector, were designed to clone the full length of human Gelsolin CDS sequences.

Forward: 5’-GTGGAATTCGCCACCATGGCTCCGCACCGCCC-3’;

Reverse: 5’-CCCTCTAGACTCGAGGGCAGCCAGCTCAGCCATGG-3’.

Primers, inducing the extra homology arms paired with the pcDNA3.1-myc-his vector, were designed to clone the full length of human GluA1 CDS sequences.

Forward: 5’-TGGCTAGTTAAGCTTGCCACCATGCAGCACATTTTTGCCTTCTT-3’;

Reverse: 5’- TGCTGGATATCTGCAGAATTCCAATCCCGTGGCTCCCAAGGGCAT-3’.

Then, the sequence and linearized pCMV-flag, pCMV-myc and pcDNA3.1-myc-his vector were recombined using Hieff Clone^®^ Plus One Step Cloning Kit (Yeasen Biotechnology (Shanghai) Co.). Next, the expression vector (pCMV-flag-DIR, pcDNA3.1-GluA1-myc-his and pcDNA3.1-Gelsolin-myc-his) and the control (pCMV-flag and pcDNA3.1-myc-his) were transfected to HEK293 cells with the help of PEI40000 reagent (Yeasen Biotechnology (Shanghai) Co.). The cells were further cultured for 48 h, and the DMEM mediums (Invitrogen) containing 10% FBS (Yeasen Biotechnology (Shanghai) Co.) were collected and centrifuged with 1000 rpm velocity for 10 min at 4□. Then, the supernatants were transferred for Western blot assay. On the other hand, the transfected cells were lysed with the HEPES lysis buffer (30 mM HEPES, 150 mM NaCl, 10 mM NaF, 1% Triton X-100, and 0.01% SDS) and centrifuged with 12000 rpm velocity for 10 min at 4□, and the supernatants were transferred for Western blotting.

### Protein purification

Gelsolin purification: pcDNA3.1-gelsolin-myc-his plasmids were transfected to HEK293 cells. After 48 h later, the cell culture medium was collected and centrifuged for 10 min with the speed of 12000 rpm. The supernatant was transferred and then incubated with Ni-NTA agarose resins (Yeasen, 20503ES60). The resins were washed with Buffer A (50 mM NaH_2_PO_4_, 300 mM NaCl, and 10 mM imidazole, PH = 8.0) before binding with supernatant. Buffer B (50 mM NaH_2_PO_4_, 300 mM NaCl, and 20 mM imidazole, PH = 8.0) was used to wash the resins after incubation. The attached proteins were eluted from resins using Buffer C (50 mM NaH_2_PO_4_, 300 mM NaCl, 250 mM imidazole, PH=8.0). The eluted solution was concentrated using Millipore Amicon® Ultra-15 and was replaced by PBS. The purified gelsolin was stored in -80□ refrigerator.

GluA1 purification: the GluA1 CDS was clone to the expression vector pGEX- 4T1-GST using the following primers.

Forward: 5’-CGGCCGCATCGTGACGCCAATTTCCCCAACAATATCCAGA-3’;

Reverse: 5’-GCAGATCGTCAGTCACAATCCCGTGGCTCCCAAGG-3’.

The pGEX-4T1-GST-GluA1 recombinant plasmid was transformed into *E. coli* BL21. Then a single clonal *E. coli* was cultured in Luria-Bertani medium (LB) medium until OD_600_ = 0.6∼0.8. Next, 0.5 mM isopropyl-β-D-thiogalactopyranoside (IPTG, Sigma) was added to induce GST-GluA1 expression under 18□ for 16 h. The medium was centrifuged with 6000 rpm for 15 min at 4□. The pellet was lysed with Lysis Buffer (20 mM Tris, 5 mM DTT, and 8 M urea, pH = 8.0) followed by sonification. After centrifugation, the supernatant was diluted to 1 M urea using Equilibrium Buffer (20 mM Tris-HCl, and 0.15 M NaCl, pH = 8.0) and then loaded onto a glutathione-agarose column previously equilibrated on Equilibrium Buffer. Wash Buffer (20 mM Tris, 1 mM EDTA, 1 M NaCl, and 0.5% Triton X-100, pH = 8.0) was supplied to wash away uncombined proteins and the GST-GluA1 was eluted using Elution Buffer (20 mM Tris-HCl, 50 mM GSH, and 0.15 M NaCl, pH = 8.0). The eluted solution was concentrated using Millipore Amicon® Ultra-15 and was replaced by PBS. The purified GST-GluA1 was stored in -80□ refrigerator.

DIR purification: the DIR CDS was clone to the expression vector SUMO-Strep using the following primers.

Forward: 5’- GAAAATTTGTACTTCCAGGGCATGGTTGCAACGGGTTCCCTG-3’; Reverse: 5’-GGATGGCTCCACGGGCTGAAAATGTTGCTGAAC-3’.

The SUMO-DIR-Strep was expressed and purified from *E. coli* BL21. The transformed *E. coli* BL21 cells were cultured in LB at 37□ until the OD_600_=0.6 ∼0.8, and then were added 0.5 mM IPTG for the induction of protein expression at 18□ for 16-18 h. Cells were harvested and resuspended in lysis buffer (50 mM Na2HPO4, 300 mM NaCl, 1% SDS, and 1 mM PMSF, pH = 8.0) and lysed by sonication. The lysate was cleared by centrifugation at 30,000 g for 30 min at 4□. The supernatant was loaded on a Streptactin Beads 4FF column (Smart-Lifesciences) pre-equilibrated with lysis buffer. The column was washed with 200 ml wash buffer (50 mM Na2HPO4 pH 8.0, and 300 mM NaCl) and eluted with elution buffer (50 mM Na2HPO4 pH 8.0, 300 mM NaCl, and 3 mM D-desthiobiotin). Then proteins were concentrated and equilibrated with 1× PBS. Finally, the protein concentration was estimated by UV at 280 nm, frozen in liquid nitrogen and stored at -80□.

### Mass spectrometry (MS)

For MS analysis, the in-gel digestion was performed by using the following protocol. The strip was performed in 1% 1, 4-dithiotreitol (DTT) in the SDS equilibration buffer (50 mM Tris-Cl (pH 8.8), 6 M urea, 30% glycerol, 2% SDS and bromophenol blue) for 15 min. This step was followed by alkylation of the free sulhydryl groups by 2.5% iodoacetamide in the SDS equilibration buffer for another 15 min in the dark at room temperature. Then the strip was cut into 18 gel sections (each section about 1.0 cm in length). Each gel section was washed three times alternately with acetonitrile and 100 mM ammonium bicarbonate. During the last wash the gel slices were incubated in 100 mM ammonium bicarbonate for 15 min at 4□. The gel slices were dried by vacuum centrifugation and allowed to swell in a 50 μl trypsin solution containing trypsin (20 μg/ml) and ammonium bicarbonate (50 mM) for 45 min at 4□. After adding another 50 μl of trypsin solution, the gel slices were kept for 20 h at 37□. The supernatant was transferred to another vial, and the gel slices were extracted for 15 min three times by 0.1% formic acid in 60% acetonitrile. The recovered peptide solutions were dried by vacuum centrifugation and desalted and cleaned using a Ziptip (Millipore, Corp., Bedford, MA).

The peptide mixtures from each section of the strip were separated by Reverse phase HPLC (RP-HPLC) followed by tandem mass analysis. RP-HPLC was performed on a surveyor LC system (Thermo Finnigan, San Jose, CA). The C18 column (RP, 180 μm × 150 mm) was obtained from Column Technology Inc. (Fremeont, CA). The pump flow was split 1:120 to achieve a column flow rate of 1.5 μl/min. Mobile phase A was 0.1% formic acid in water, and mobile phase B was 0.1% formic acid in acetonitrile. The tryptic peptide mixtures were eluted using a gradient of 2-98% B over 180 min.

The MS was performed on a LTQ linear ion trap mass spectrometer (Thermo Finnigan, San Jose, CA) equipped with an electrospray interface and operated in positive ion mode. The capillary temperature was set to 170□ and the spray voltage was at 3.4 kV. Normalized collision energy was at 35%. Automatic gain control was used to obtain maximal signal of each scan. The mass spectrometer was set so that one full MS scan was followed by ten MS/MS scans on the 10 most intense ions. Dynamic exclusion was set at repeat count 2, repeat duration 30 s, and exclusion duration 90 s.

The acquired MS/MS spectra was searched against the IPI human database using BioWorks 3.0 software (Thermo Finnigan) on an 8 node Dell PowerEdge 2650 cluster. An accepted SEQUEST result had a ΔCn score of at least 0.1 (regardless of charge state), a value known for high confidence in a SEQUEST search. All output results were combined by using a homemade software named Build Summary to delete the redundant data. To make sure that the MS/MS spectrum was of good quality with fragment ions clearly above baseline noise, we referred to the parameters reported in previous studies and applied stricter criteria for the peptide identification. Peptides were validated after meeting the following criteria. The SEQUEST cross-correlation score must be ≥ 1.9 for a + 1 tryptic peptide, ≥ 2.2 for a + 2 tryptic peptide and ≥ 3.75 for a + 3 tryptic peptide. In addition, ΔCn cutoff values were ≥ 0.1 and the SP rank of the peptides ≤ 4.

### ThT fluorescence assay

ThT binding assays were modified from the previously reported article. Briefly, Monomeric Aβ_42_(1 mM) was dissolved with DMSO and diluted to a final concentration of 10□μM with ThT fluorescence assay buffer (50□mM sodium phosphate buffer (pH = 7.4), 50□mM NaCl, 1□μM ThT and 0.01% sodium azide). Samples were added to a 96-well black plate and incubated at 37□°C without shaking. Real-time ThT fluorescence measurements were taken using a Tecan Spark microplate reader (Tecan). The fluorescence values were measured every 10 - 20□min for 3 - 4□h at excitation and emission wavelengths of 440□nm and 480□nm, respectively.

### Brain slice preparation

Adult mice were anaesthetized and decapitated, then the brains were quickly removed and placed in ice-cold artificial cerebrospinal fluid (ACSF) containing the following compounds (in mM): 117 NaCl, 3.6 KCl, 1.2 NaH_2_PO_4_·2H_2_O, 2.5 CaCl_2_·2H_2_O, 1.2 MgCl_2_·6H_2_O, 25 NaHCO_3_, and 11 glucose. The ice-cold ACSF had a pH of 7.4 when bubbled with 95% O_2_ and 5% CO_2_. The brains were glued onto the stage of a Leica VT1200S vibratome and 350 μm-thick transverse hippocampal slices were cut. The slices were incubated for at least 1 h in oxygenated (95% O_2_ and 5% CO_2_) ACSF at room temperature before recording.

### Electrophysiology

For spontaneous excitatory postsynaptic current (sEPSC), the slices were transferred to a submerged chamber that was mounted on the stage of an upright microscope (Olympus, BX51WIF), and perfused with oxygenated (95% O_2_ and 5% CO_2_) ACSF. The patch pipettes (2.5 – 3.5 MΩ) were fabricated from borosilicate glass tubes (WPI, 1B150F-3) using a horizontal puller. The pipettes solutions contained (mM): 135 K-gluconate, 0.5 CaCl_2_, 2 MgCl_2_, 5 KCl, 5 EGTA, 5 HEPES, and 5 D-glucose (pH adjusted to 7.3 with KOH). Spontaneously occurring excitatory postsynaptic currents (sEPSCs) of hippocampal CA1 region recordings were performed using an amplifier (Multiclamp 700B, Molecular Devices) under the voltage clamp, with cells clamped at -70 mV. Signals were filtered at 2 kHz. All currents were sampled using a Digidata 1,550B interface. Traces were recorded in pClamp10.5 software (Molecular Devices).

For the long-term potentials, hippocampal slices were placed on submersion recording chambers perfused with oxygenated (95% O_2_ and 5% CO_2_) ACSF (30 ± 0.5□, 2 ml/min). fEPSPs responses were evoked by stimulating the Schaffer collateral with 0.2 ms pulses delivered through customized double barrel glass electrodes (Sutter Instrument Co.) and recording from stratum radiatum of area CA1 using glass pipettes filled with ACSF. The stimulation intensity was systematically increased to determine the maximal fEPSP slope and then adjusted to half maximal fEPSPs slope. fEPSPs were recorded, filtered at 0 – 1000 Hz, digitized and stored at 5K sampling rate using customized program in Spike hound. fEPSPs slope were analysed using customized Matlab code. Initial slopes of fEPSPs were expressed as percentages of baseline averages. After 30 min stable baseline recorded, LTP was induced by 4 trains of theta-burst stimulation (TBS) at 0.1 Hz. Each train consists of 10 bursts (4 pulses at 100 Hz per burst) repeated at 5 Hz.

### Transmission Electron Microscope (TEM)

For TEM, two hundred microliters of 5 μM monomeric Aβ42 alone, 5 μM monomeric Aβ42 and 1 μM IR peptides added with or without 0.1 μM gelsolin were incubated overnight in RIPA lysis buffer at 4□. Then, the samples were centrifugated (15,000g for 5 min at 4□) to collect the pellets and resuspended with twenty microliters of RIPA lysis buffer. The TEM samples were bound to carbon-coated grids and stained with 2% uranyl formate. Finally, images were collected using TEM (Thermo Fisher Scientific) operated at 120 kV.

### Proximity ligation assay (PLA)

To test interaction of DIR, gelsolin and Aβ in situ, paraffin section of human brain tissues was provided by Human Brain Bank, Chinese Academy of Medical Sciences & Peking Union Medical College, Beijing, China. Slides were blocked and then incubated in a humidity chamber for 60 min at 37°C. Then the antibodies recognized DIR, gelsolin and Aβ were supplied to stain the slides overnight at 4°C. On the next day the slides were incubated with PLUS and MINUS PLA probes (1:5; Sigma; DUO92001 and DUO92005) for 1 h at 37°C followed by ligation with 1X Ligation Buffer (1:40; Sigma; DUO92008) and the additional incubation for 30 min at 37°C. Moreover, the slides were incubated with Amplification-Polymerase solution (1:80; Sigma; DUO92008) for 100 min at 37°C. Finally, mount the slides with a cover slip using a minimal volume of Duolink In Situ Mounting Medium with DAPI (Sigma; DUO82040). Leica SP8 microscope was used for imaging.

### Hypoxia treatment

Neural stem cells and differentiated neurons were placed in a chamber system (Stemcell^TM^ technologies) with the hypoxic condition of 1% O_2_ and 21% O_2_ as control condition for 24 h. Then, the cells were lysed with the Lysis Buffer (100 mM Tris, 150 mM NaCl, 8 M urea and protease inhibitors, PH = 7.5) followed by sonification, and the lysates were centrifuged with 12000 rpm velocity for 10 min at 4□, and the supernatants were measured with Western blotting.

Mice were accommodated in laboratory mouse cage continues supplied with hypoxic condition (8% O_2_) through air inlet hole for 8 h. Oxygen analyzer (Shanghai Precision Instrument Co., Ltd) were detected in air outlet hole ensuring the hypoxic condition of the cage.

### Animal model of cerebral ischemia

The ischemia model based on middle cerebral artery occlusion (MACO) surgery, was modified according to the published paper^56^. Briefly, mice were anesthetized with isoflurane and then a skin lesion in the middle of the neck was cut. After gently separating the muscle, the common carotid artery (CCA) and its branches were exposed under a stereo dissecting microscope. Then the CCA was gently ligated by silk sutures to slower the blood flux while the distal external carotid artery (ECA) was tightly ligated to block the flux. Next, a small incision of the ECA between the branch point and ligation was generated and the tip rounded suture (Beijing Cinotech Co.) was inserted along the ECA into the internal carotid artery (ICA) until to occlude the origin of middle cerebral artery (MCA). Furthermore, tightly ligation between incision and branch point of ECA was performed while the suture was slowly withdrawn from CCA. Finally, the surgical region was closed.

### *In vivo* injection

The DIR-Intron was released to the bilateral hippocampus of mouse brain by osmotic pumps system (ALZET®, 1002). Briefly, pumps were implanted subcutaneously to direct deliver the DIR-Intron peptide or the scrambled peptide (dissolved in PBS) to the injection site via the brain infusion kit (ALZET®) for two weeks. mDIR siRNAs (SS-GCGUUGUUCCAGUCUGCUATT; AS-UAGCAGACUGGAACAACGCTT) were purchased from Guangzhou RiboBio Co., Ltd. In total, 15 nmol unit siRNAs were injected to bilateral hippocampi of WT or APPPS1 mice for 3 times with 3 days intervals between each injection. Injection sites were the CA1 regions of bilateral hippocampi (from bregma, siRNAs were injected ± 2.0 mm lateral, -2.25 mm caudal, -1.85 mm ventral).

### *In vivo* two-photon imaging

Three to 5-month-old mice were anesthetized with 1.5% isoflurane and fixed to a stereotaxic frame with custom-built stage-mounted ear bars. Then a cranial window was implanted on the head of mouse. A 1.5 cm incision was made between the ears, and the scalp was reflected to expose the skull followed by making a circular craniotomy (5 – 6 mm diameter) using a high-speed drill and a dissecting microscope for gross visualization. Next, a glass-made coverslip was attached to the skull. Then, anti-DIR antibodies were conjugated with FITC (Abcam, ab188285) under the product instructions. Finally, a two-photon microscope turned to 920-nm and 1045-nm wavelength lasers were used to acquire images after intravenous injection of Dextran-Texas Red (Thermo Fisher Scientific, D1830) and DIR antibodies.

### Behavior phenotyping

*Y-maze:* The animals were placed in one arm end of three identical arms (40 × 40 × 40 cm with the 120° angle between arms) and given free access to the arms for a 10-min session. The entry of mice into each arm was video-recorded, and the sequence and number of arms entered were analyzed.

*Novel object recognition test:* Adult mice were placed in and familiar with an open box (40 × 40 × 40 cm). 12∼24 h later, two identical objects were set to the box and mice were allowed to explore for 10 min. To test for object recognition after training, one of the objects was replaced with a novel object, and the animals were entered in the apparatus again for 10 min. The time spent on exploring each of the objects was measured. All movement were recorded by a camera.

*Morris water maze:* Morris water maze was performed as previously described, but with minor modifications. The experimental procedures consisted of 2 phases: training phase and test trials. The training phase consisted of 4 trials daily with different entries (30-min interval) for 5∼6 days. A hidden platform was set 0.5 cm under the water. Mice were allowed to swim freely to find the platform for 1 min. If not completed within the specified period, the correct routes would be offered by operators. Mice were allowed to stay on the platform for 30 s before being picked up. The test trials were performed after training, mice were placed in water and the platform was removed. The escape latency and the time spent in each quadrant were recorded by a camera and were analyzed with Etho Vision XT 14 software.

*Open-field test:* Mice were placed in an open field (45 × 45 cm) and recorded the exploratory locomotor activity in a 30 min period by a Digiscan apparatus (Accuscan Electronics). The moving distance and velocity were analyzed.

A detailed account of the methods and statistical analyses used in this study is provided in the supplementary materials.

### Statistical Analysis

The data are presented as mean ± SEM. Sample number (n) values are indicated in figures, figure legends or results section. Two groups were compared by a two-tailed, unpaired Student’s t test. Comparisons between two groups with multiple time were performed by a two-way ANOVA with Bonferroni’s post hoc test.

Statistical analysis was performed using PRISM (GraphPad Software). Differences were considered significant at p < 0.05 (*p < 0.05, **p < 0.01, ***p < 0.001).

## Acknowledgements

These studies were funded by QuietD. The authors thank the patients in clinic studies. We thank Dr. Y.Q. Li for help in the histological data, Dr. J.S. Guan and Dr. K.W. He for LTP recording, Dr. Z.N. Zhang for providing human ES cells, Dr. J.J. Chen and X.Y. Zen for plasmid construction and cell line culture, Hao Feng (National Center for Protein Science Shanghai, China) for animal management.

## Contributions

K.C.L. conceived and designed the project. X.Z. and Y.M. provided advise for the project. H.X.S., P.Y., J.P., L.D., Y.C.C., S.Z. and B.C. performed plasmid construction, cell line culture, immunostaining and IP. Z.L., J.W.L., X.F.M. and Y.L. performed the electrophysiological and behavioral experiments. L.C. provided the clinic data. H.B.W. and Y.Q.Z. advised for experiment and manuscript. K.C.L., X.Z., H.X.S., Z.L. and P.Y. performed statistical analyses and made the figures. K.C.L., X.Z., Y.M. and H.X.S. prepared the manuscript.

Present address of X.Z. and B.C.: Guangdong Institute of Intelligence Science and Technology, Hengqin 519031, Zhuhai, Guangdong, China.

## Supplementary information

### Supplementary figure legend

**Fig. S1:**
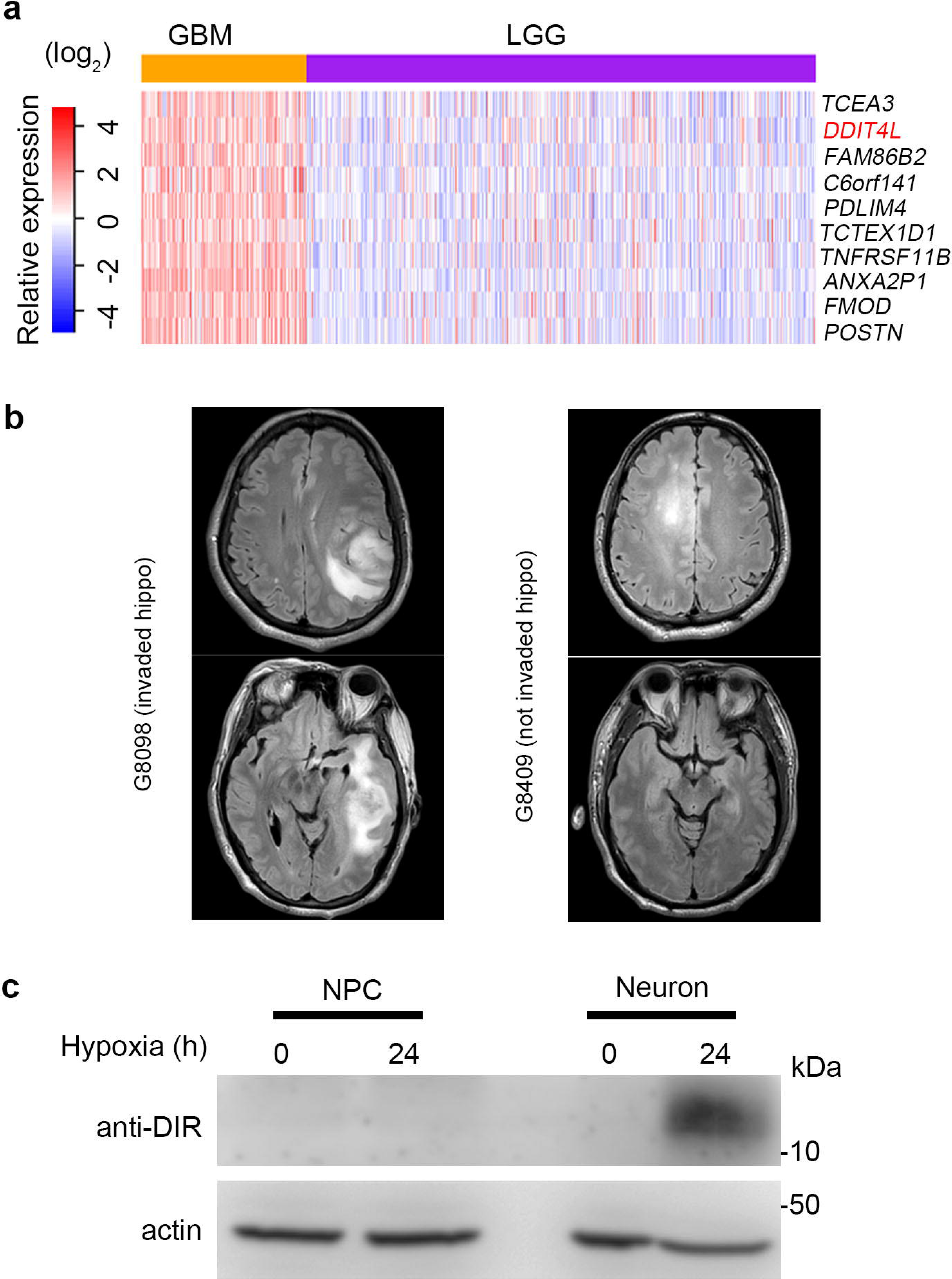
DDIT4L expression in aggressive glioma, and DIR in human stem cell-derived neurons under hypoxia and AD patients. a. Top 10 upregulated genes were identified in the GBM&LGG comparative analysis from the TCGA database. b. Magnetic resonance imaging of the patient’s brain with the GBM invaded the hippocampus (hippo, G8098 in Fig. 1f) or the brain with GBM not in the hippo (G8409 in Fig. 1f). c. The neurons derived from human stem cells, not the neural precursor cells (NPC), enhanced the expression of DIR under hypoxia (1% O_2_, 24 h).

**Fig. S2:**
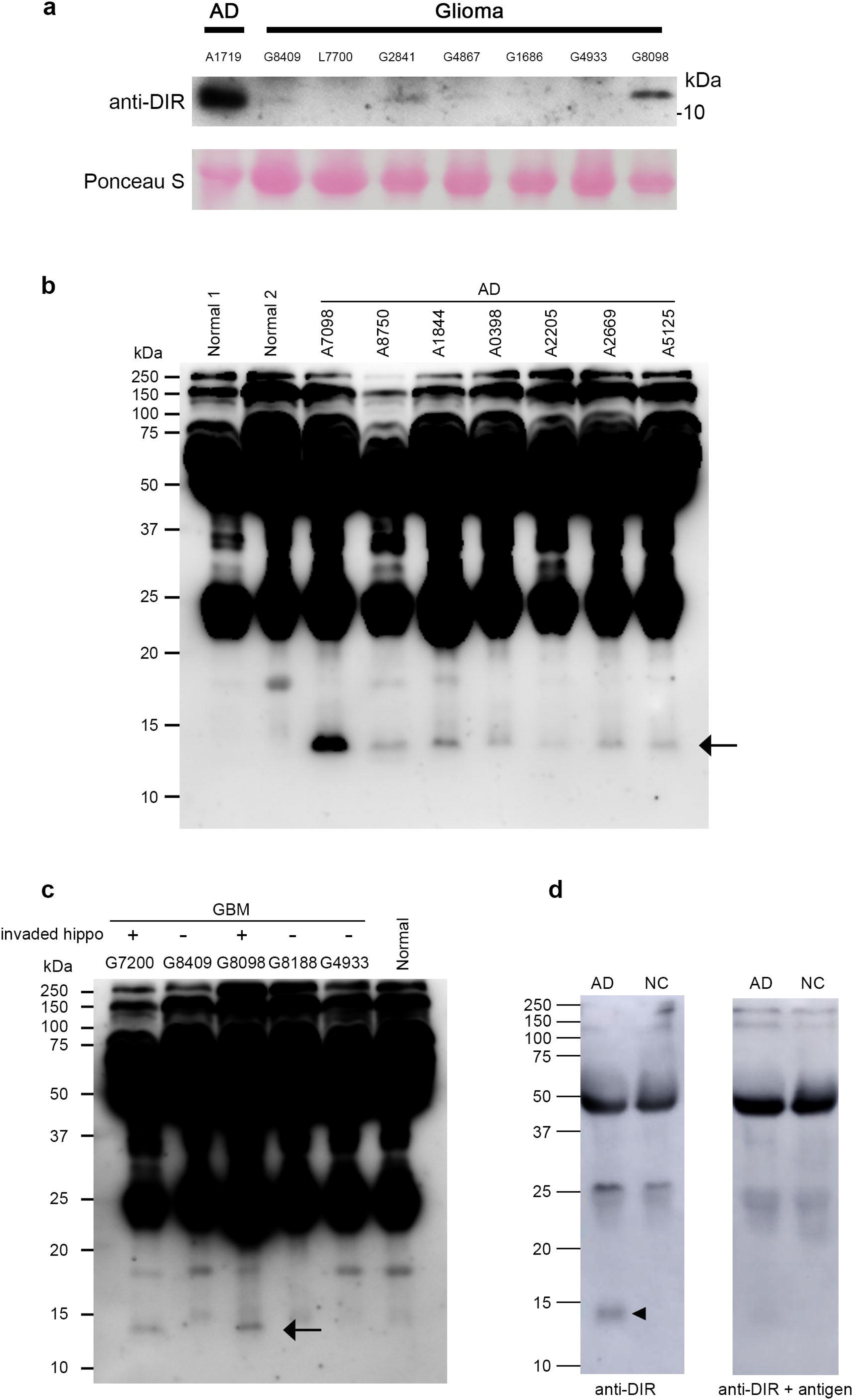
The expression of DIR in plasma. a. In the plasma of AD and glioma patients, the intensity of DIR immunoblot of AD patients was higher than that of the glioma patients. b. The full lanes of DIR in plasma of AD patients, related to Fig. 1g. c. The full lanes of DIR in plasma of GBM patients, related to Fig. 1f. d. The DIR antibody absorption verification in plasma of normal control and AD patients.

**Fig. S3:**
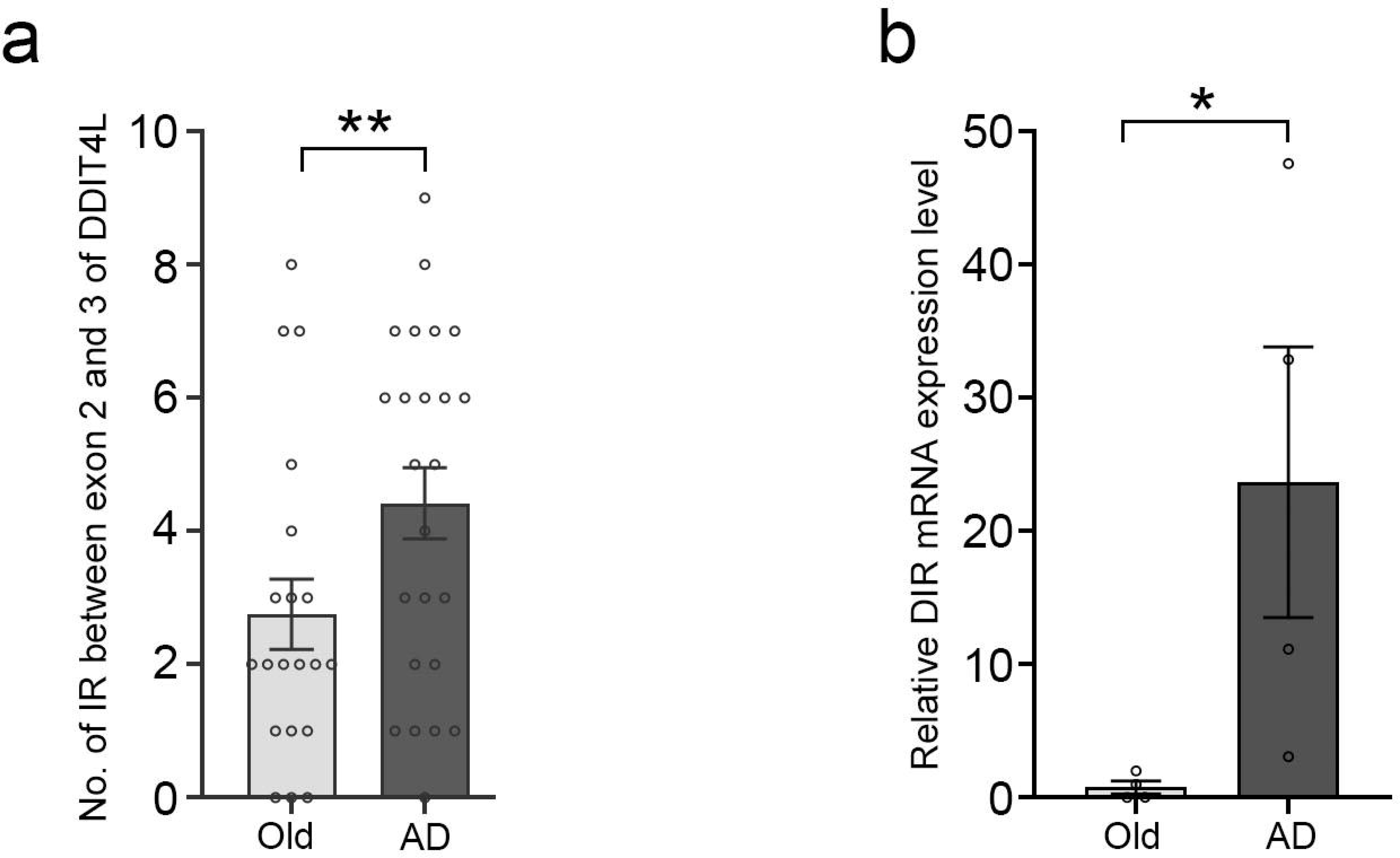
The intron retention of DIR in normal control and AD patients. a. The IR expression analysis of DDIT4L using IRfinder. b. The relative expression of DIR mRNA in normal old people and AD patients.

**Fig. S4:**
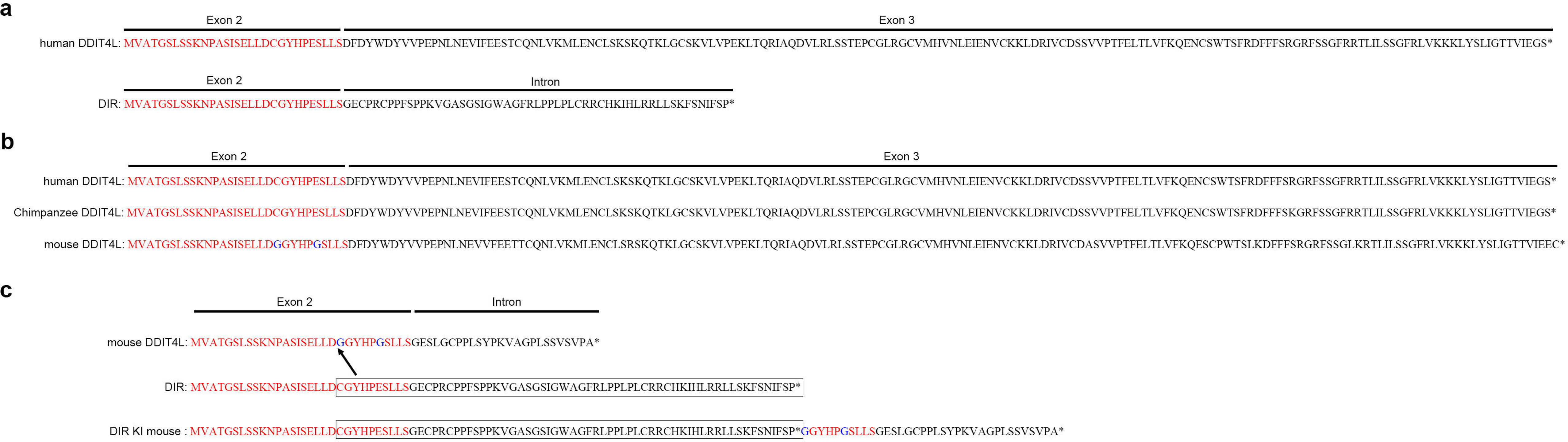
Human DDIT4L and DIR protein sequences, species difference and human DIR knock-in sequence in mouse. a. The amino acid (AA) sequences of human DDIT4L and DIR. b. Comparison among human, chimpanzee and mouse DDIT4L sequences. c. The AA sequences of constructed DIR expressed in the DIR-KI mice.

**Fig. S5:**
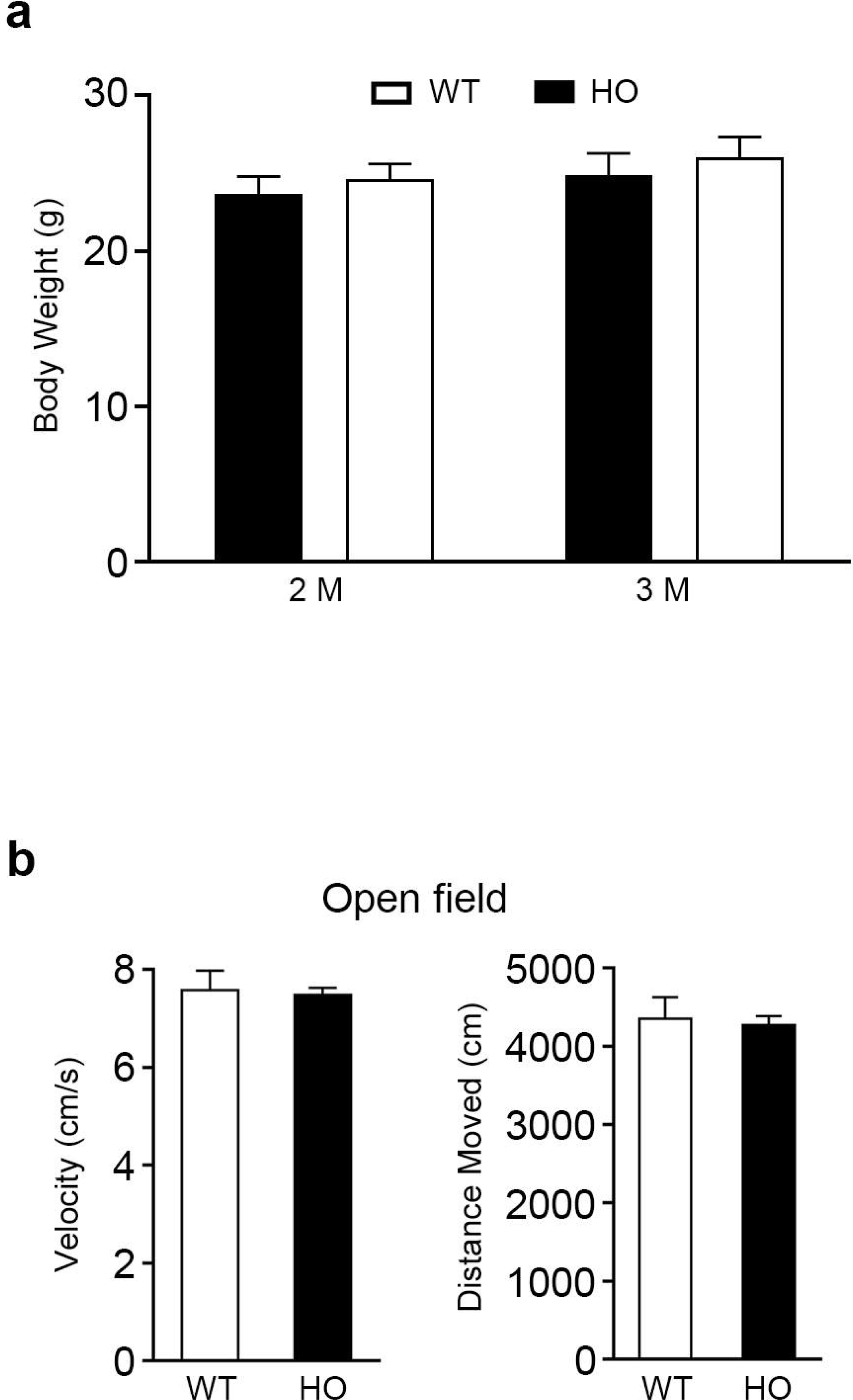
Hippocampal expression of DIR didn’t affect growth and motor ability of homozygous DIR-KI mice. a. The body weight of the homozygous DIR-KI mice (n = 8) was not different from that of WT mice (n = 8). b. The open field test showed no difference in the moving distance and velocity between WT mice (n = 10) and the homozygous DIR-KI mice (n = 17).

**Fig. S6:**
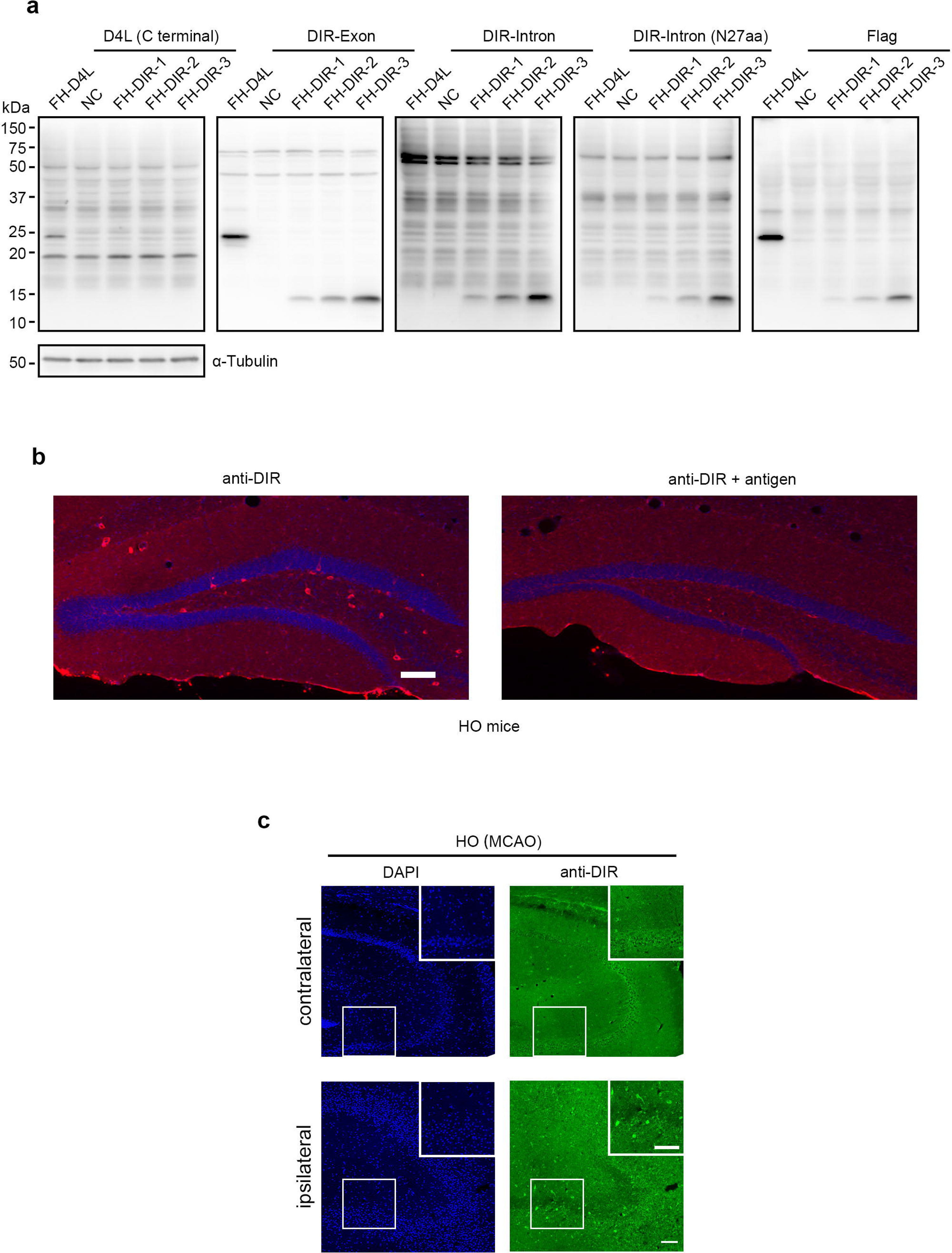
The antibody verification. a. The immunoblotting was used to test the specificity of anti-DDIT4L, anti-DIR antibodies. b. The DIR antibody absorption verification in homozygous DIR-KI mice. Scale bar = 50 mm. c. The Immunohistochemistry showed the DIR expression was apparently increased in the ipsilateral hippocampus in MCAO hypoxic-ischemic brain injury model of homozygous DIR-KI mice (n = 4). Scale bar = 100 mm.

**Fig. S7:**
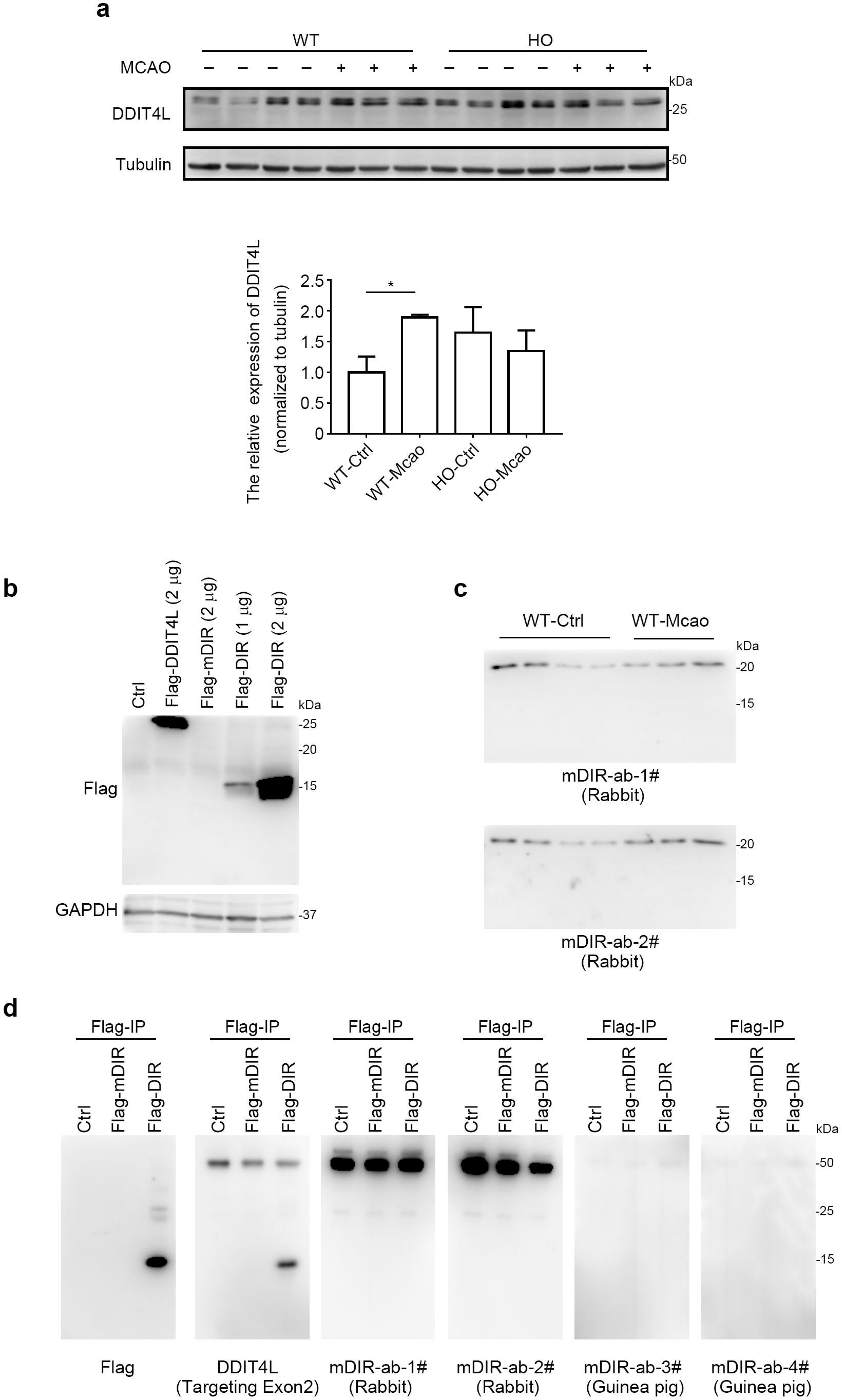
The detection of mDDIT4L and mDIR. a. The detection of mDDIT4L in hippocampus of normal control and MCAO model between WT and DIR-KI mice. *, p < 0.05. b. The flag-mDIR plasmid was not expressed in HEK293 cells. c. The mDIR was not detected in normal control and MCAO model of WT mice. d. The flag-mDIR was not detected after flag antibody immunoprecipitation.

**Fig. S8:**
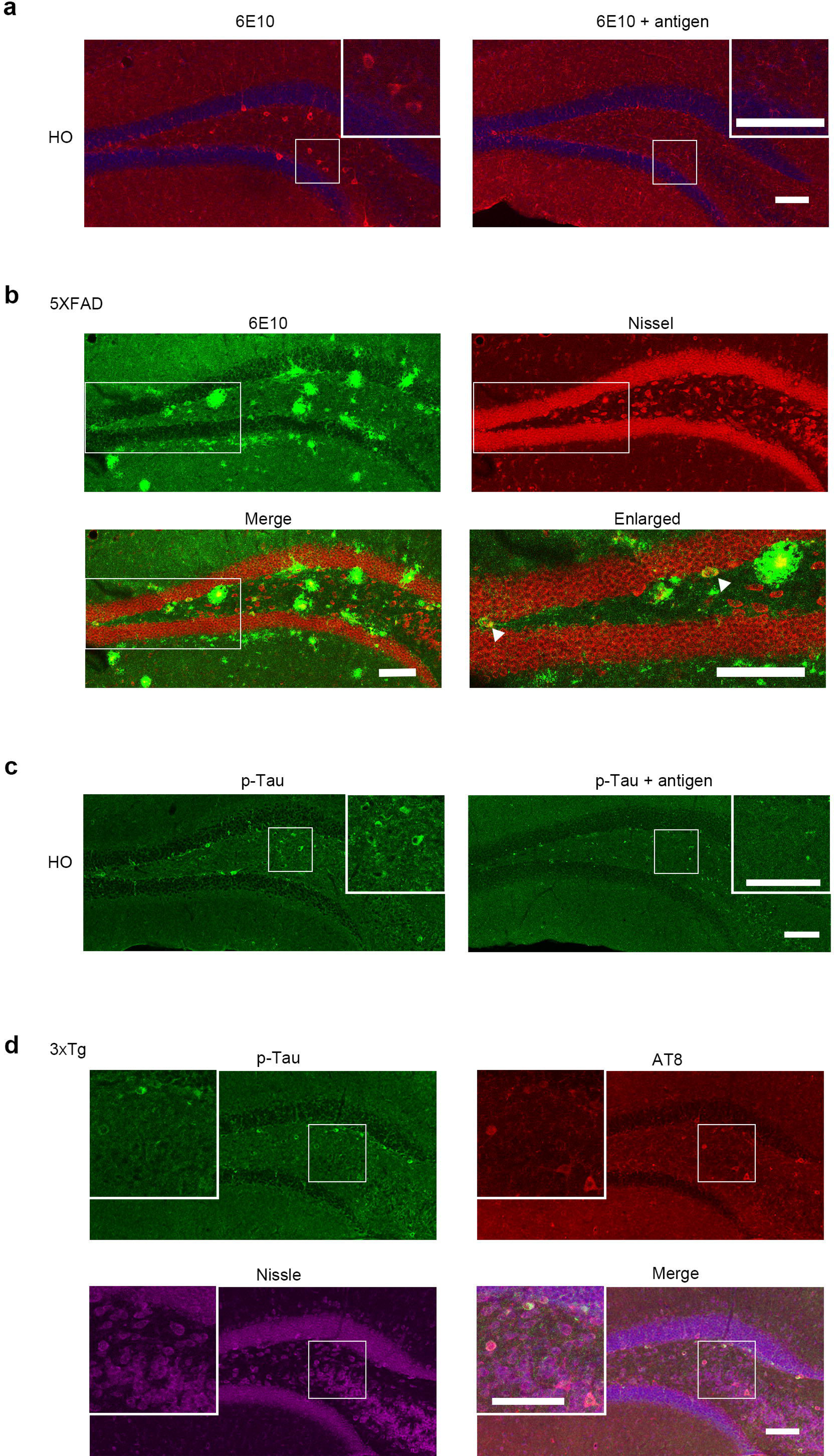
The verification of Aβ and p-Tau antibodies. a. The antigen absorption verification of Aβ antibody (6E10) in DIR-KI mice. Scale bar = 100 μm. b. The 6E10 was used to detect Aβ in 5XFAD mice. Scale bar = 100 μm. c. The antigen absorption verification of p-Tau antibody in DIR-KI mice. Scale bar = 100 μm. d. The pTau antibody was used to detect Aβ in 3XTg mice. Scale bar = 100 μm.

**Fig. S9:**
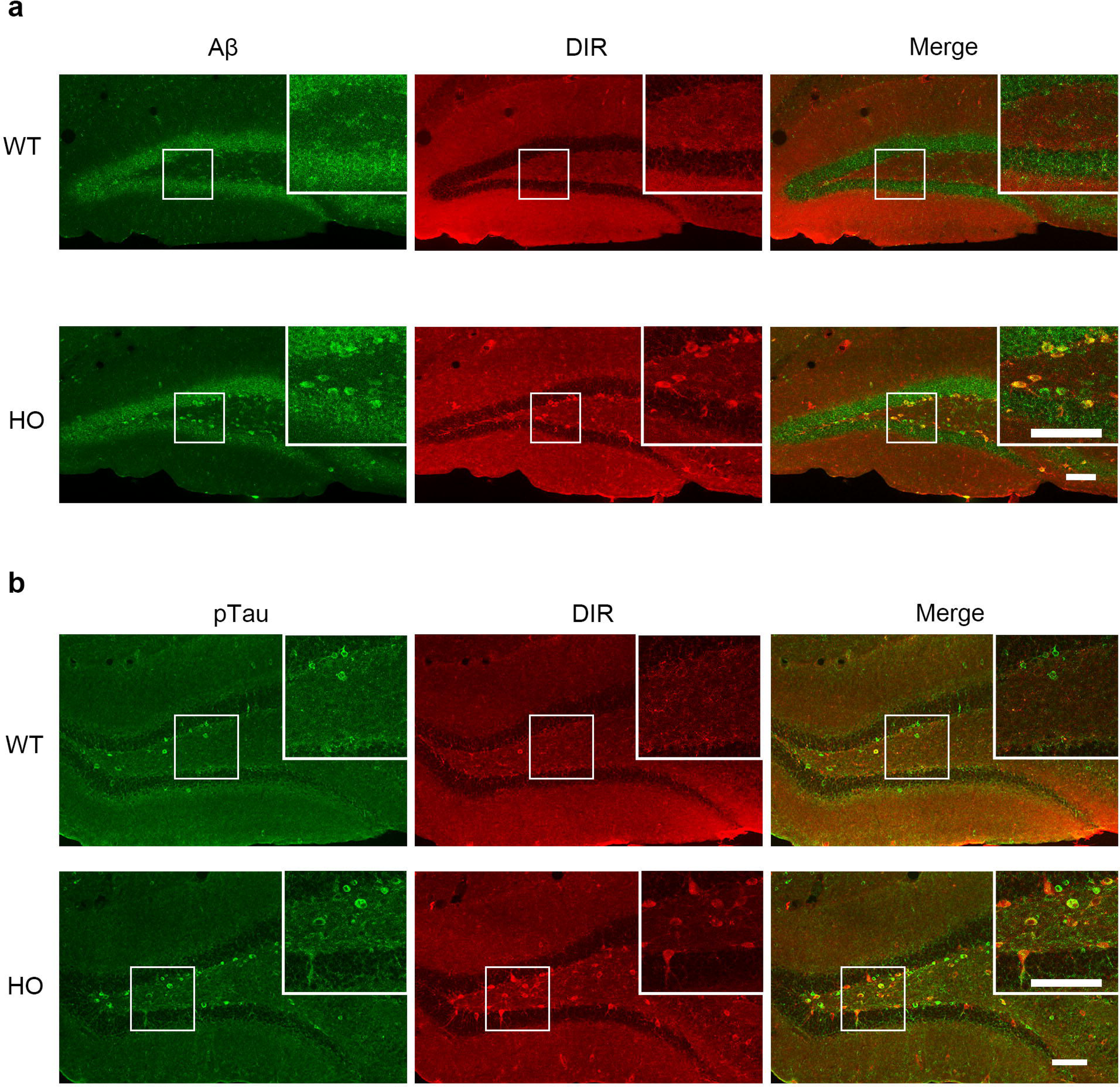
DIR colocalized with Aβ and p-Tau. a. The colocalization of DIR and Aβ in hippocampus of DIR-KI mice. Scale bar = 100 μm. b. The colocalization of DIR and p-Tau in hippocampus of DIR-KI mice. Scale bar = 100 μm.

**Fig. S10:**
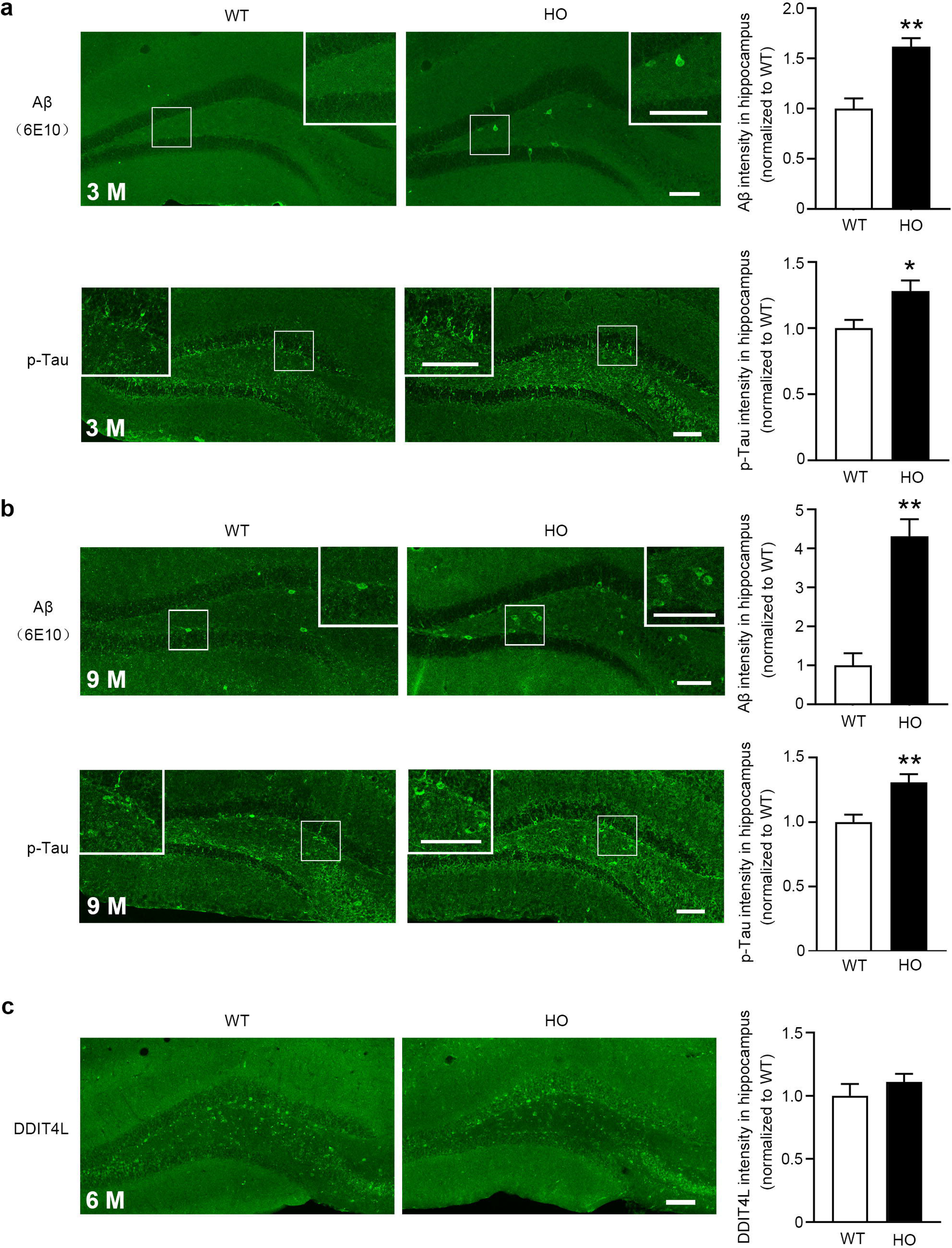
Aβ- or p-Tau expression in 3, 9-month-old DIR-KI mice and DDIT4L expression in 3-month-old DIR-KI mice. a. The expression of Aβ- or p-Tau was upregulated in the hippocampal dentate gyrus in the 3-month-old homozygous DIR-KI mice. Scale bar = 100 μm. *, p < 0.05. **, p < 0.01. b. The expression of Aβ- or p-Tau was upregulated in the hippocampal dentate gyrus in the 9-month-old homozygous DIR-KI mice. Scale bar = 100 μm. **, p < 0.01. c. The expression of DDIT4L have no change in the hippocampal dentate gyrus in the WT and homozygous DIR-KI mice. Scale bar = 100 μm.

**Fig. S11:**
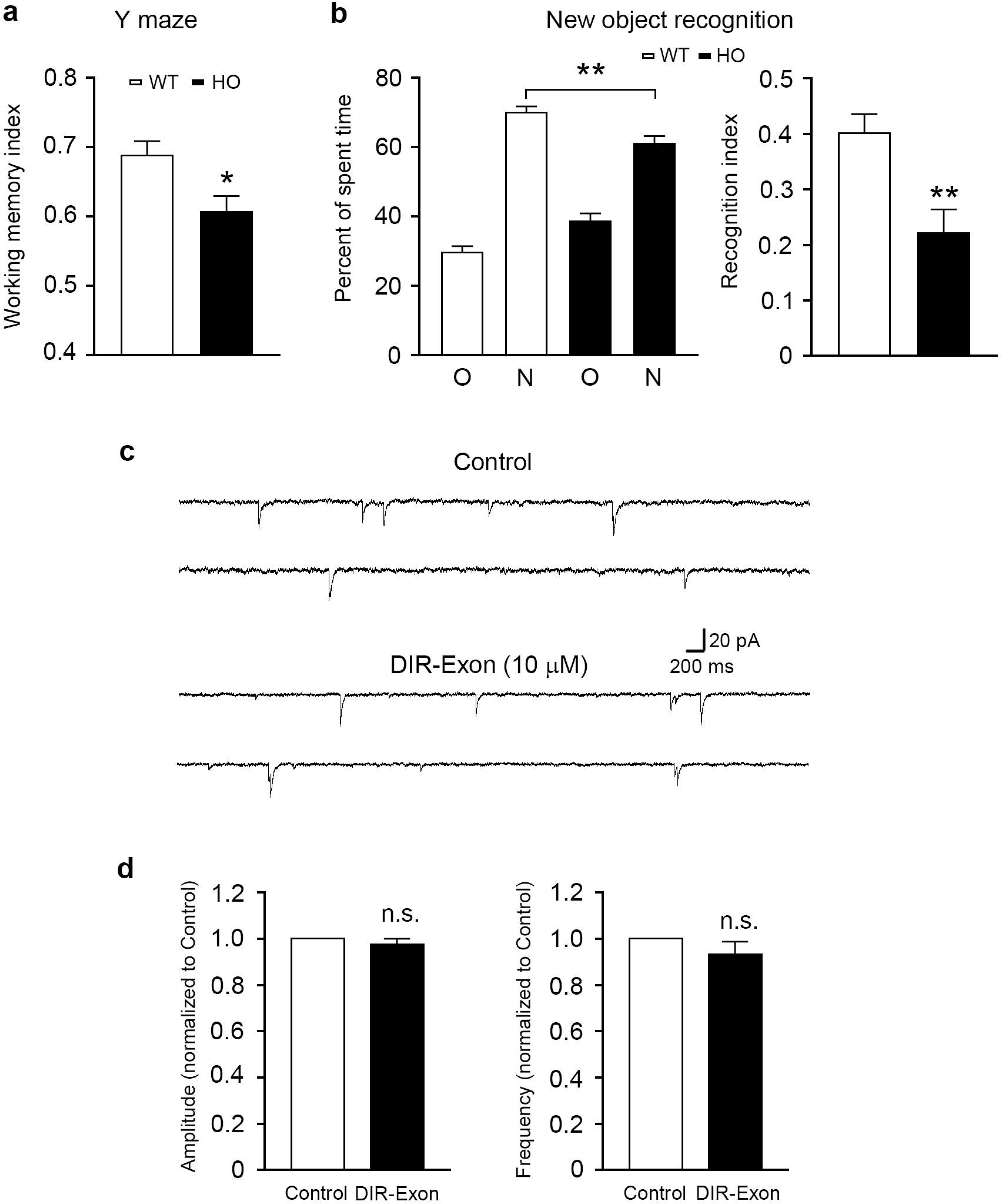
Working memory and novel object recognition in DIR-KI mice. a. The Y maze alternative test showed that the working memory was impaired in the homozygous DIR-KI mice (HO, n = 22), as compared with that of wild-type mice (WT, n = 20). *, p < 0.05. b. The capacity of novel object recognition was reduced in the homozygous DIR-KI mice (n = 17) compared with that of WT mice (n = 19). **, p < 0.01. c. The sEPSC of hippocampal neurons was not altered by applying DIR-Exon in the brain slices prepared from WT mice. d. The amplitude and frequency of sEPSC in hippocampal neurons were not changed by DIR-Exon (10 μM, n = 9).

**Fig. S12:**
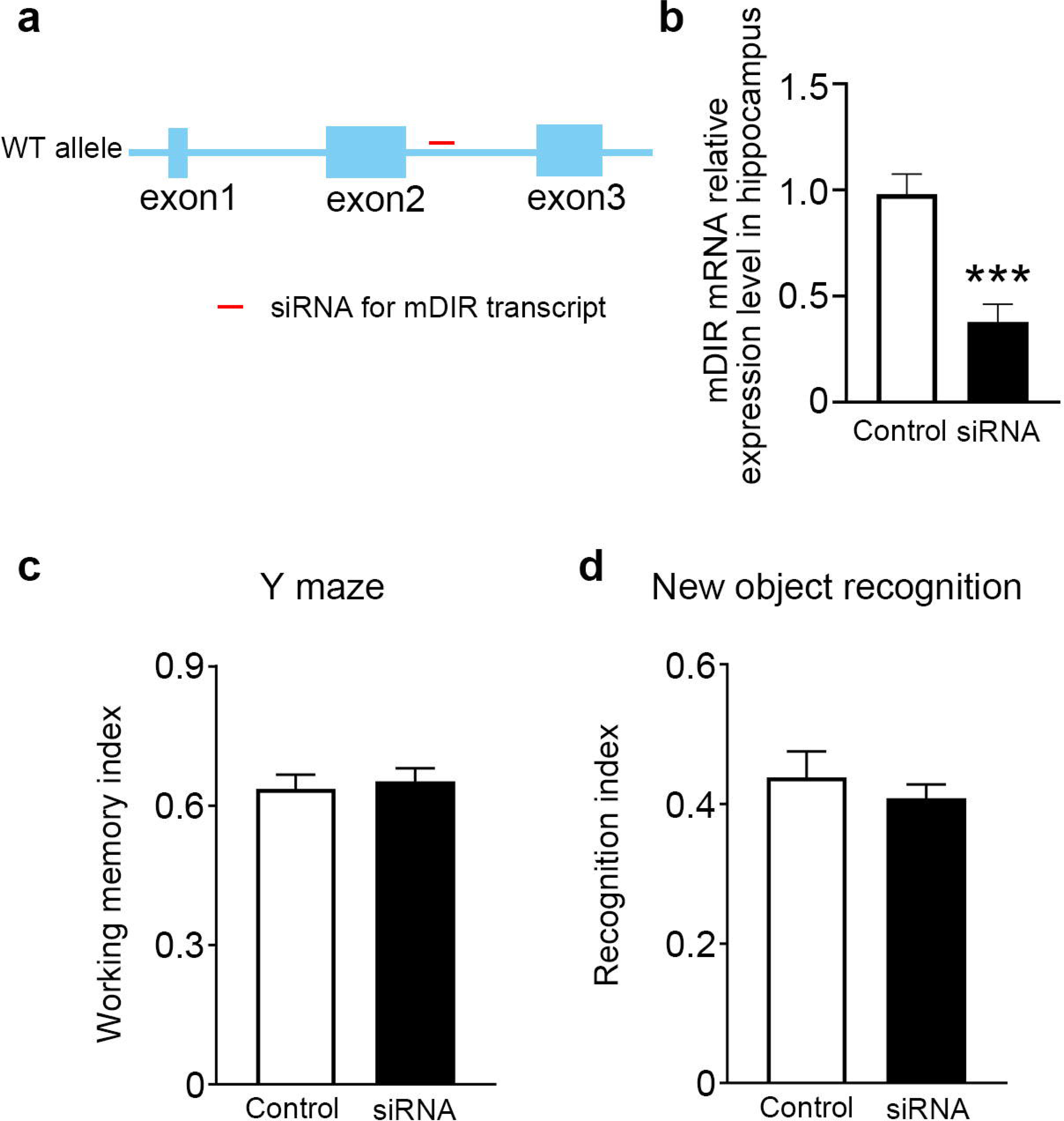
No effects of mDIR on working memory and novel object recognition. a. Presentation the action site of siRNA for mDIR transcript in the WT allele. b. The siRNA apparently reduced mDIR mRNA level (n = 5). ***, p < 0.001. c. The Y maze alternative test showed that the working memory had no change between the control (n = 9 mice) and siRNA (n = 10 mice) groups. d. The capacity of novel object recognition had no change between the control (n = 9 mice) and siRNA (n = 10 mice) groups.

**Fig. S13:**
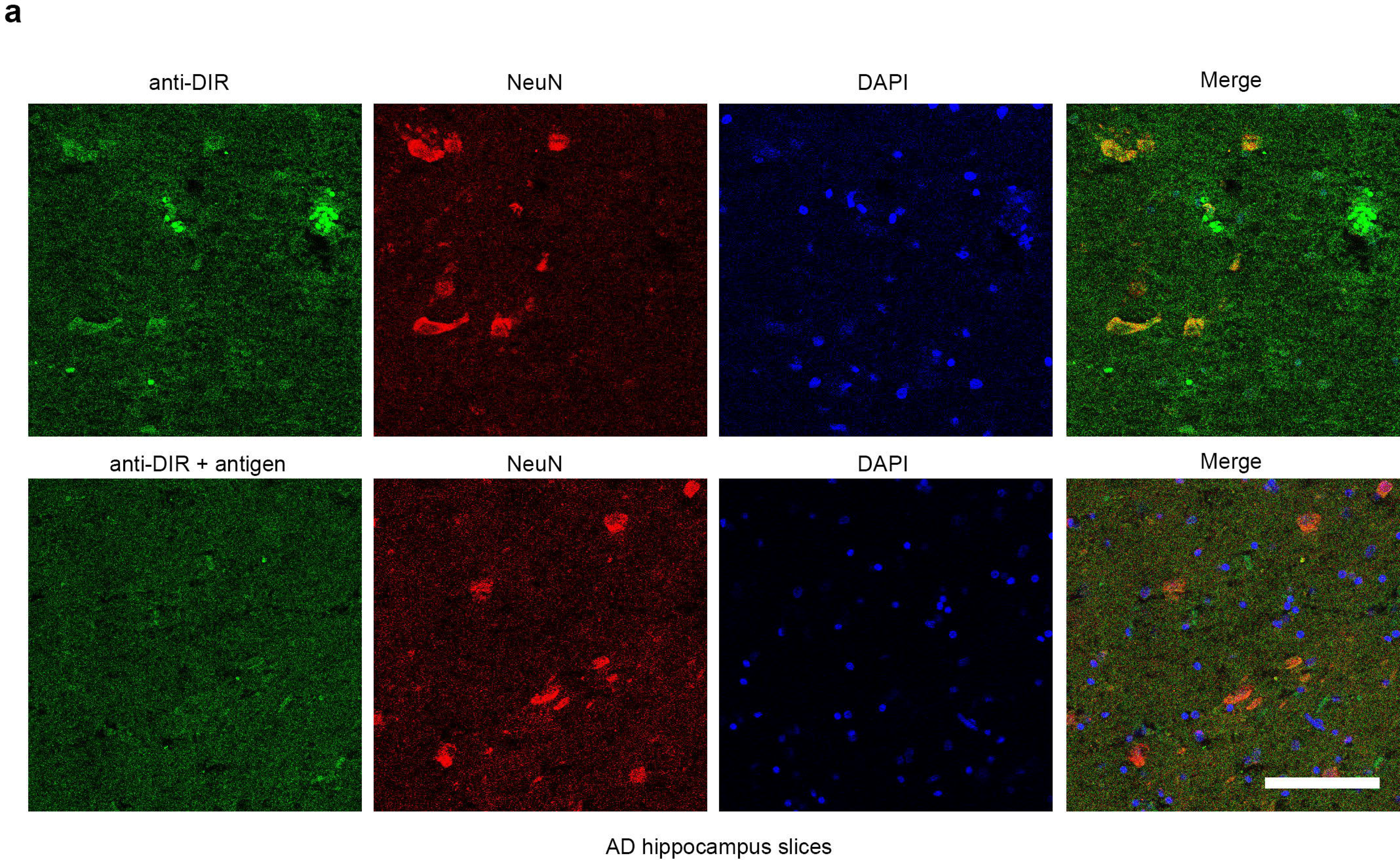
DIR expression in hippocampus in AD patient. a. The antigen absorption verification of DIR antibody in hippocampus slices of AD patients. Scale bar = 100 μm.

**Fig. S14:**
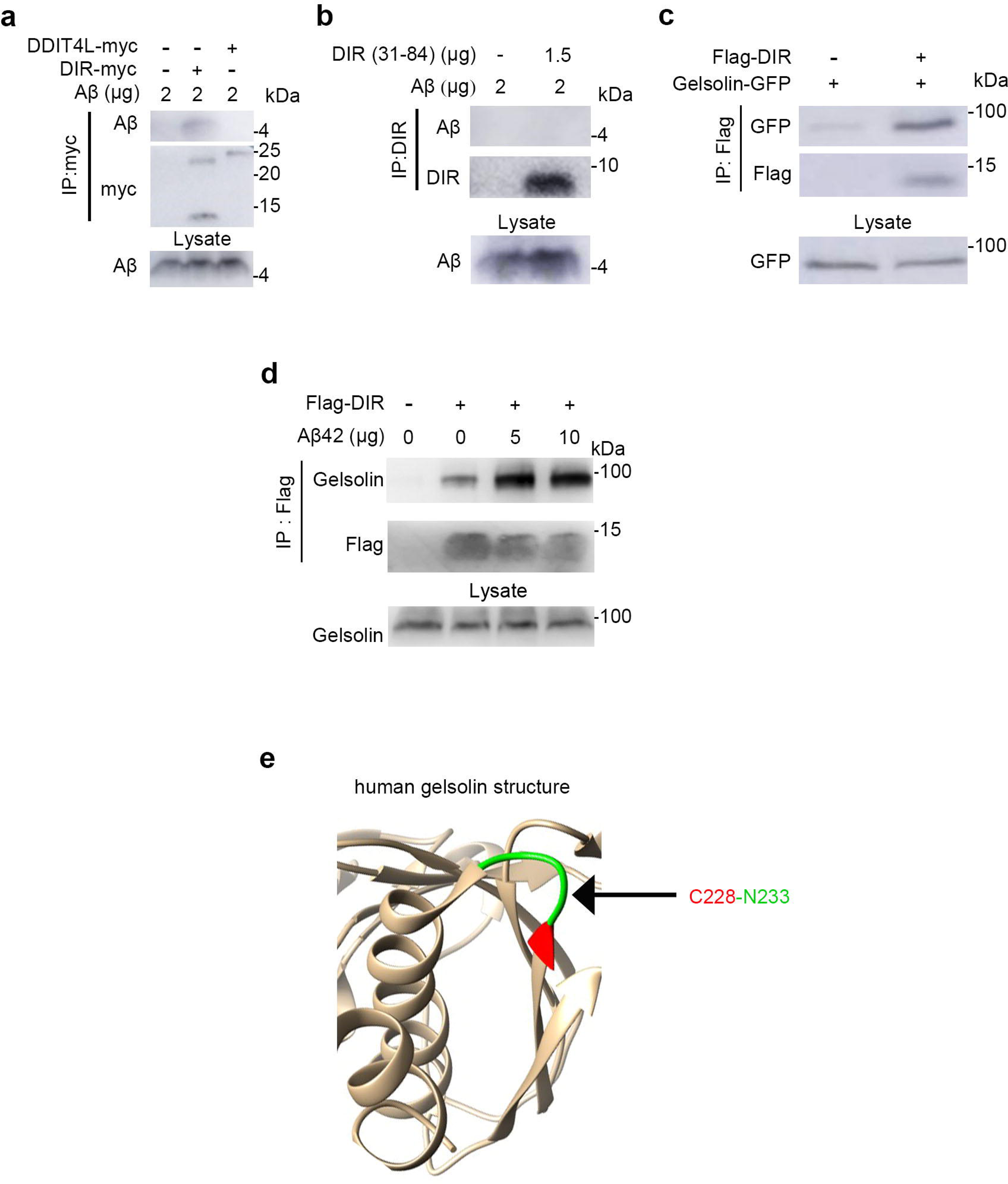
DIR directly bind to gelsolin but not Aβ. a. In the mixture of lysate of HEK293T cells co-transfected plasmids expressing myc-DIR or DDIT4L-myc with additions of synthetic Aβ42, Aβ was found in the proteins precipitated with myc antibodies in the lysate of expressing DIR-myc (n = 3). b. Co-IP with two synthetic proteins, DIR (31-84) and Aβ42, showed no direct interaction between DIR (31-84) and Aβ42 (n = 3). c. In the lysate of HEK293T cells co-transfected with plasmids expressing Flag-DIR and gelsolin-GFP, gelsolin-GFP was found in the proteins precipitated with Flag antibodies (n = 3). d. In the lysate of HEK293T cells transfected with the plasmid expressing Flag-DIR with additions of different doses of synthetic Aβ42, gelsolin was found in the proteins precipitated with Flag antibodies, and increased binding level in the group with high doses of Aβ42 (5, 10 μg, n = 3). e. The C228-N233 motif in gelsolin (Gelsolin^C228-N233^) could be the docking site for DIR. C228 in gelsolin could be the key point for the interaction between DIR and gelsolin.

**Fig. S15:**
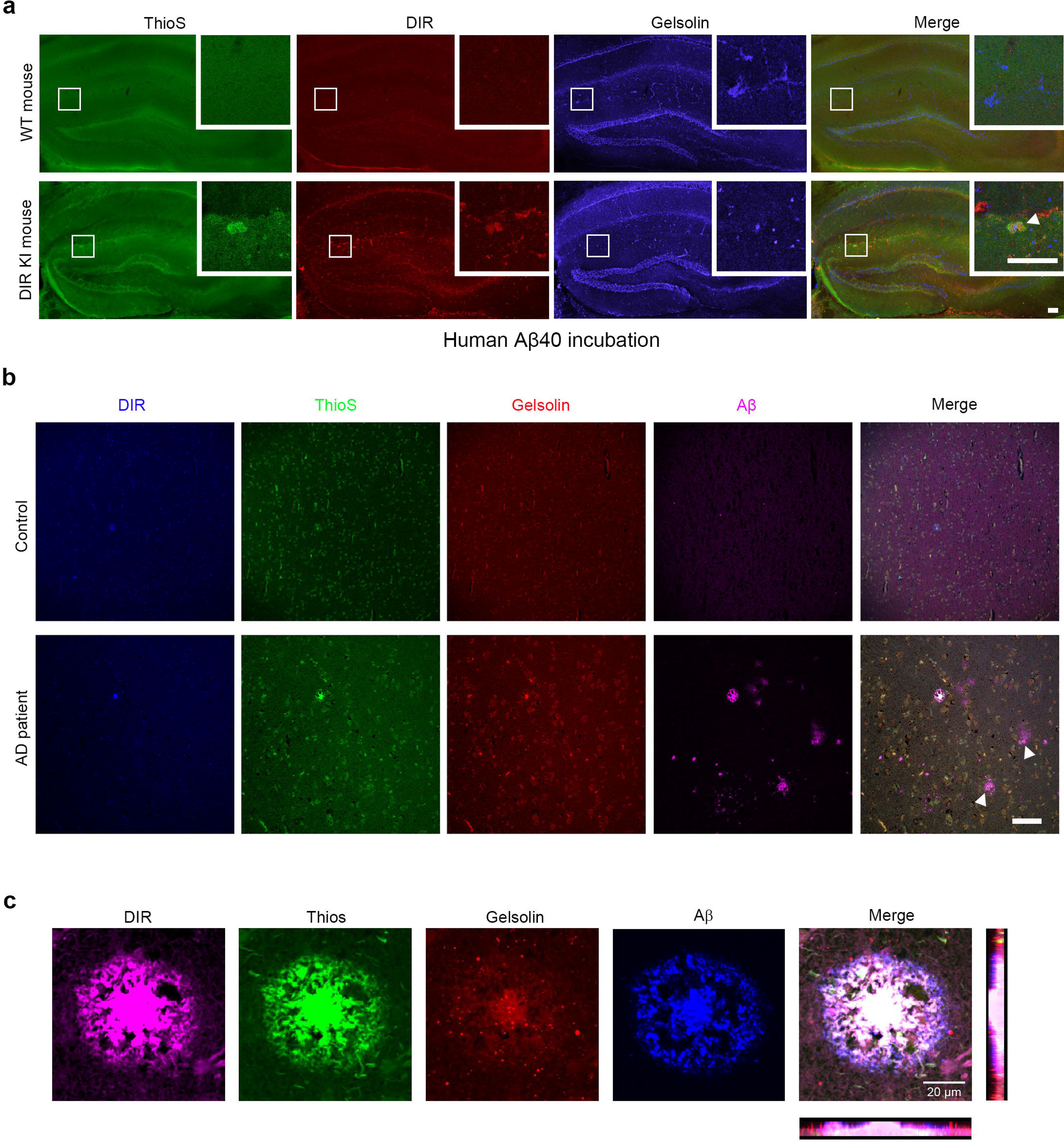
The co-localization of DIR, Aβ and gelsolin. a. In the hippocampus slices of the homogenous DIR-KI mice (n = 3), but not that of WT mice (n = 3), incubated with the synthetic human Aβ40 (100 μM) for 6 h, thioflavine S-positive plaque (arrow) appeared and co-existed with DIR and gelsolin. Scale bar = 100 μm. b. In AD patients (n = 3), but not control individuals (n = 3), the Aβ-positive, thioflavine S-negative diffuse plaque (arrows) was not co-localized with DIR and gelsolin in the hippocampal DG area. Scale bar = 50 μm. c. The confocal image of DIR, Aβ, gelsolin and ThioS in hippocampus of AD patients. Scale bar = 20 μm.

**Fig. S16:**
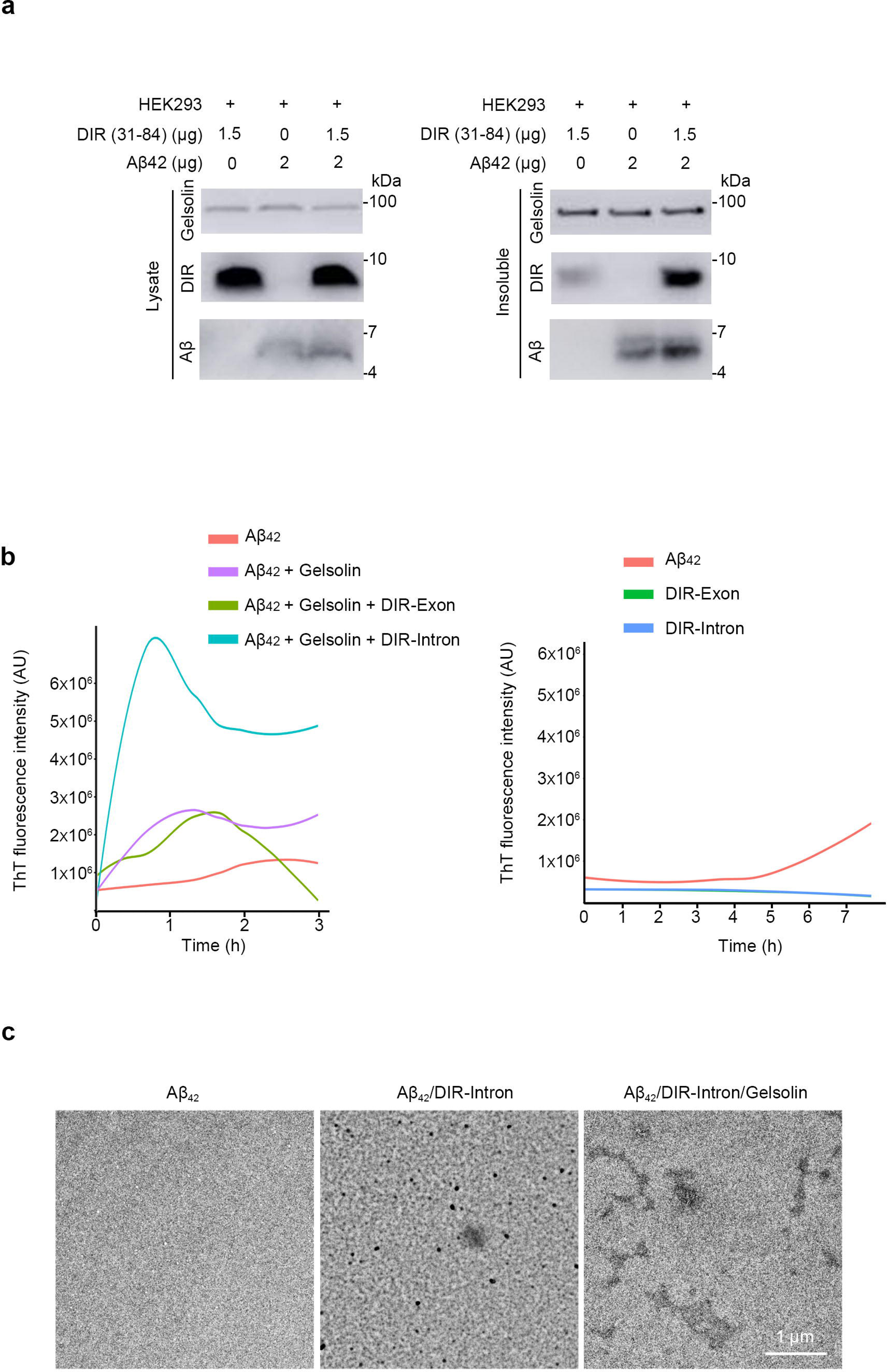
DIR induces Aβ deposition via gelsolin. a. In the mixture of lysate of HEK293T cells with additions of synthetic Aβ42 or DIR (31-84), DIR (31-84) induced Aβ42 insolubilization via gelsolin. The DIR decreased and Aβ disappeared in the proteins precipitated with gelsolin antibodies (n = 3). b. The results of the ThT fluorescence assay following Aβ42, gelsolin, DIR-Exon and DIR Intron incubation separated or combined. c. The results of electron microscope assay following Aβ42, Aβ42/DIR-Intron and Aβ42/gelsolin/DIR-Intron incubation.

**Fig. S17:**
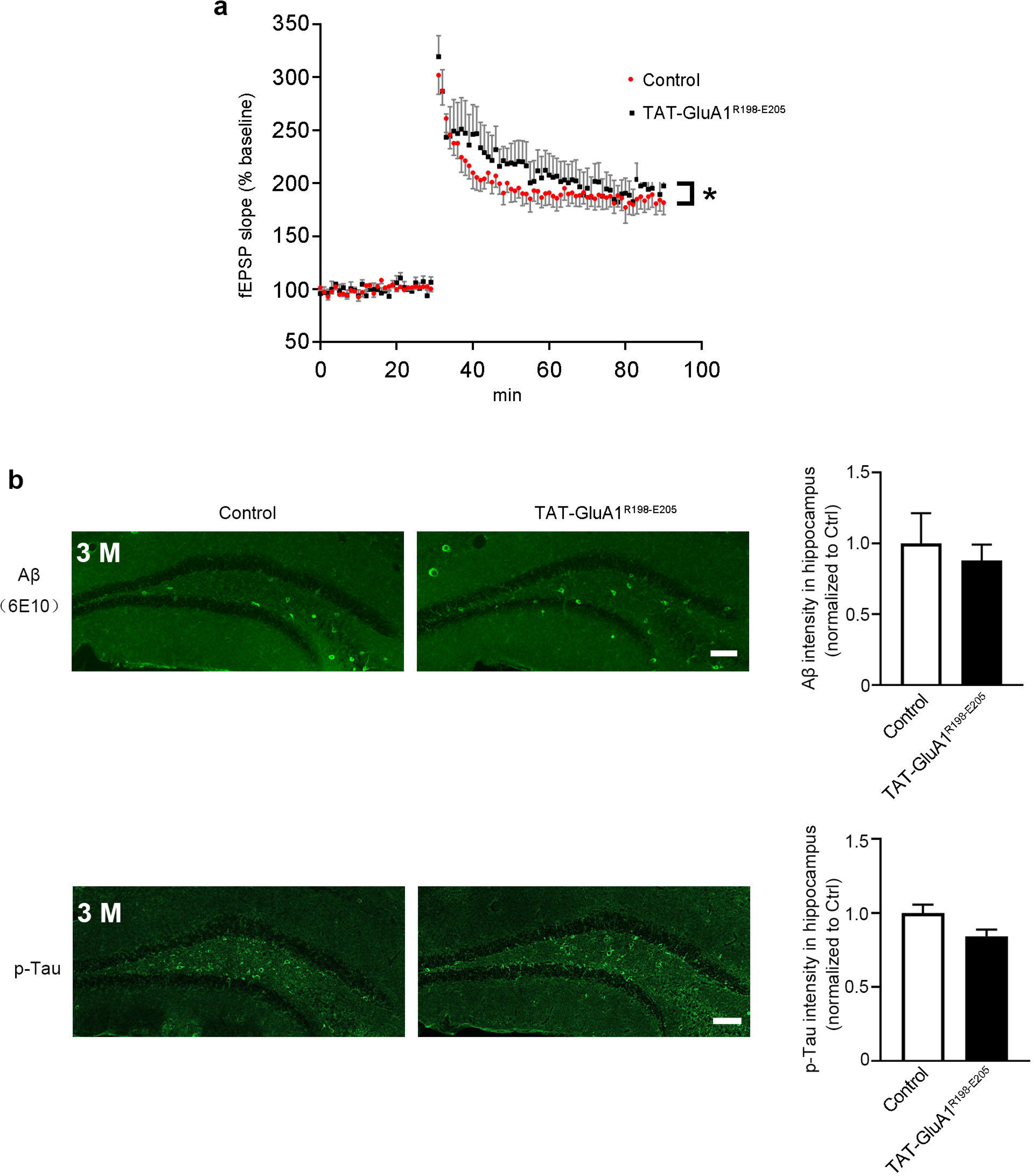
TAT-GluA1^R198-E205^ enhanced hippocampal LTP in the homozygous DIR-KI mice. a. The LTP in the homozygous DIR-KI mice was apparently increased after the TAT-GluA1^R198-E205^ treatment, as compared with control treatment (n = 5 mice per group). *, p < 0.05. b. The expression of Aβ- or p-Tau had no change in the hippocampal dentate gyrus in the 3-month-old homozygous DIR-KI mice after the TAT-GluA1^R198-E205^ treatment. Scale bar = 100 μm.

**Fig. S18:**
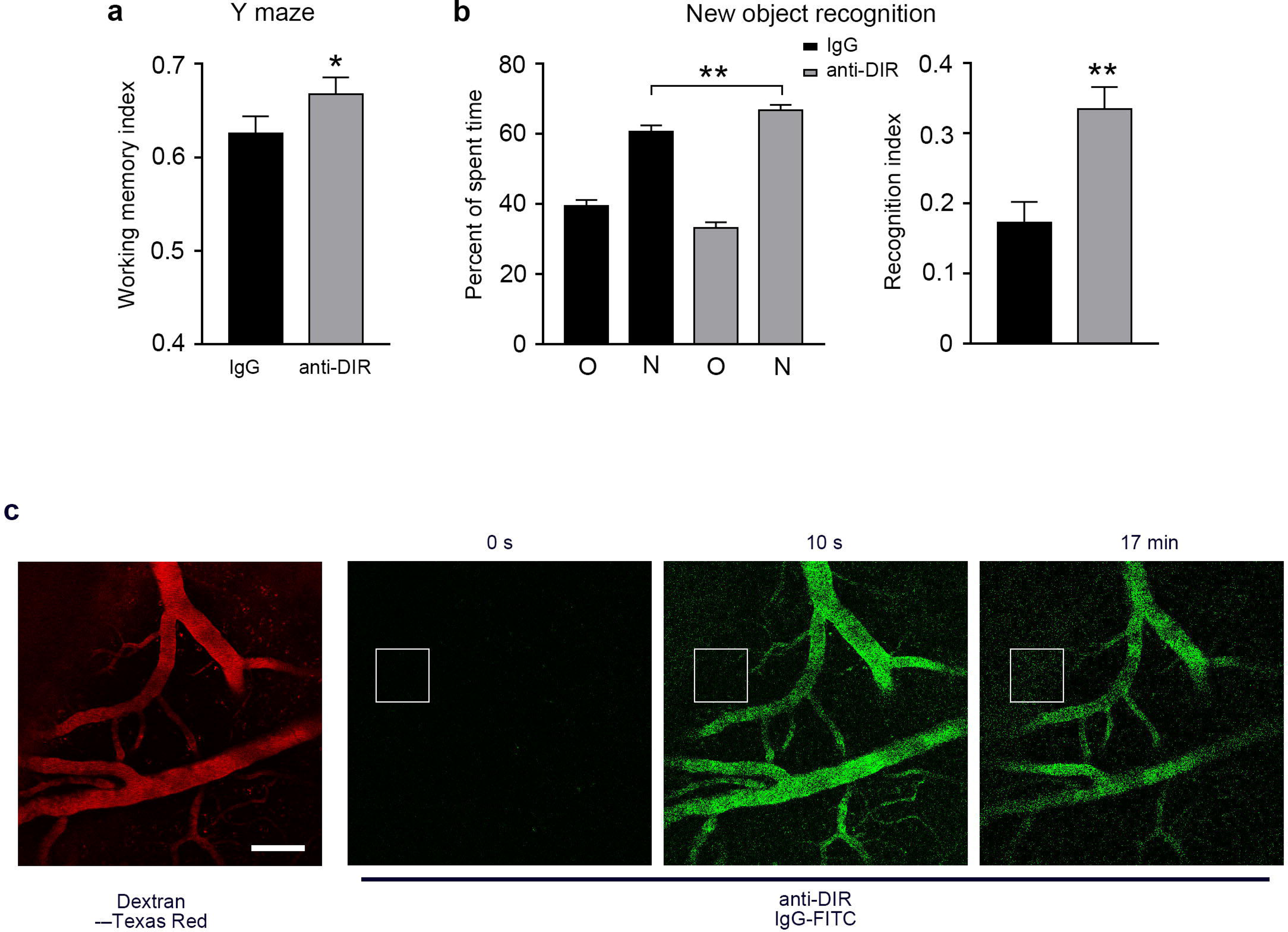
The DIR mAb improves the working memory and novel object recognition in the homozygous DIR-KI mice and could pass through the blood-brain barrier. a. The Y maze alternative test showed that the working memory was reversed after the DIR mAb treatment in the homozygous DIR-KI mice (n = 17), as compared with the mice treated with IgG (n = 13). *, p < 0.05. b. The capacity of novel object recognition was reversed by the DIR mAb treatment in the homozygous DIR-KI mice (n = 14), as compared with the mice treated with IgG (n = 11). **, p < 0.01. c. The mouse brain two-photon imaging after intravenous injection of FITC conjugated DIR antibody and Dextran-Texas Red showed that DIR antibody crossed the blood-brain barrier into brain tissue (n = 3 mice).

**Supplementary table 1:**
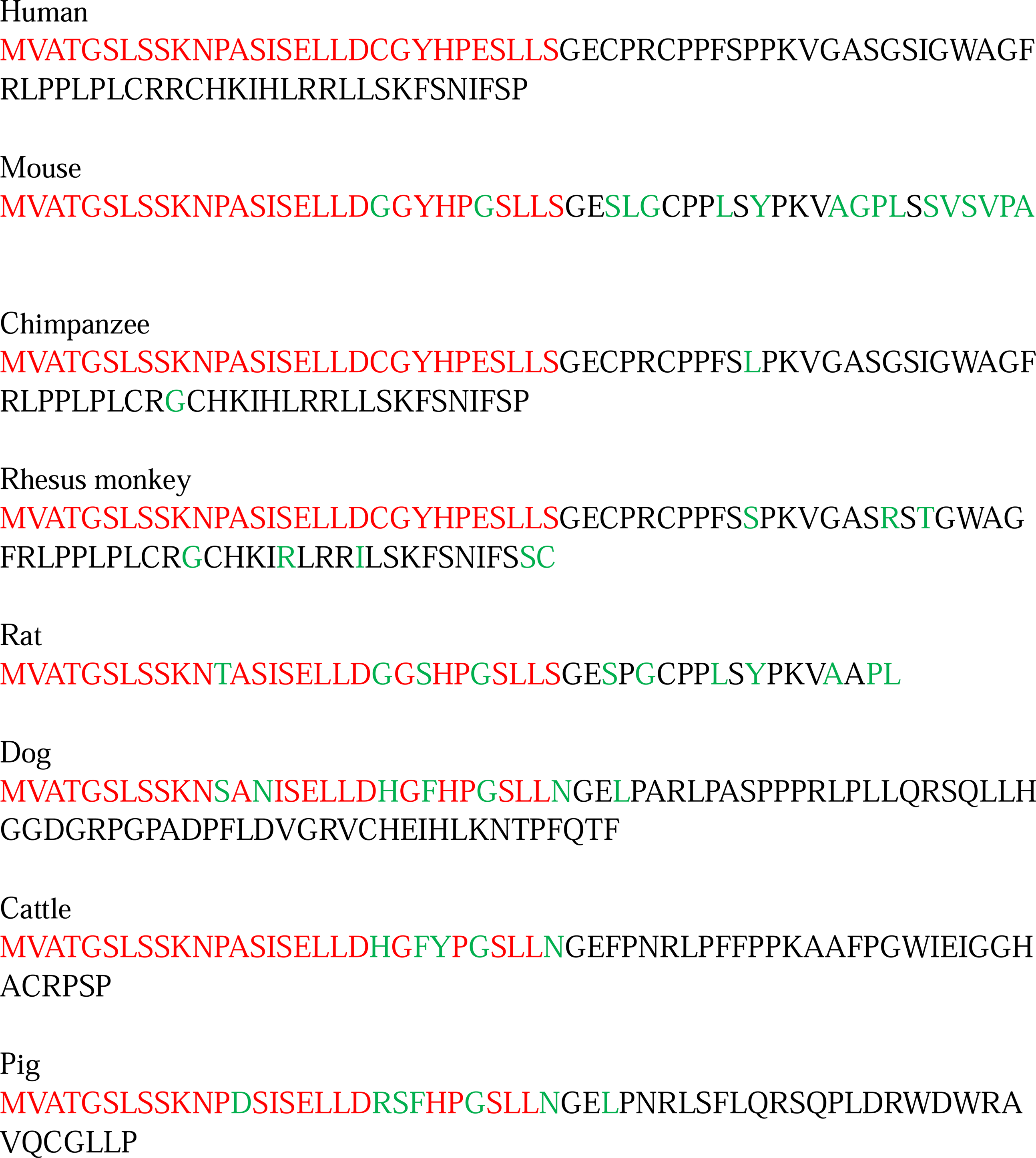
DIR in different species.

**Supplementary table 2.**
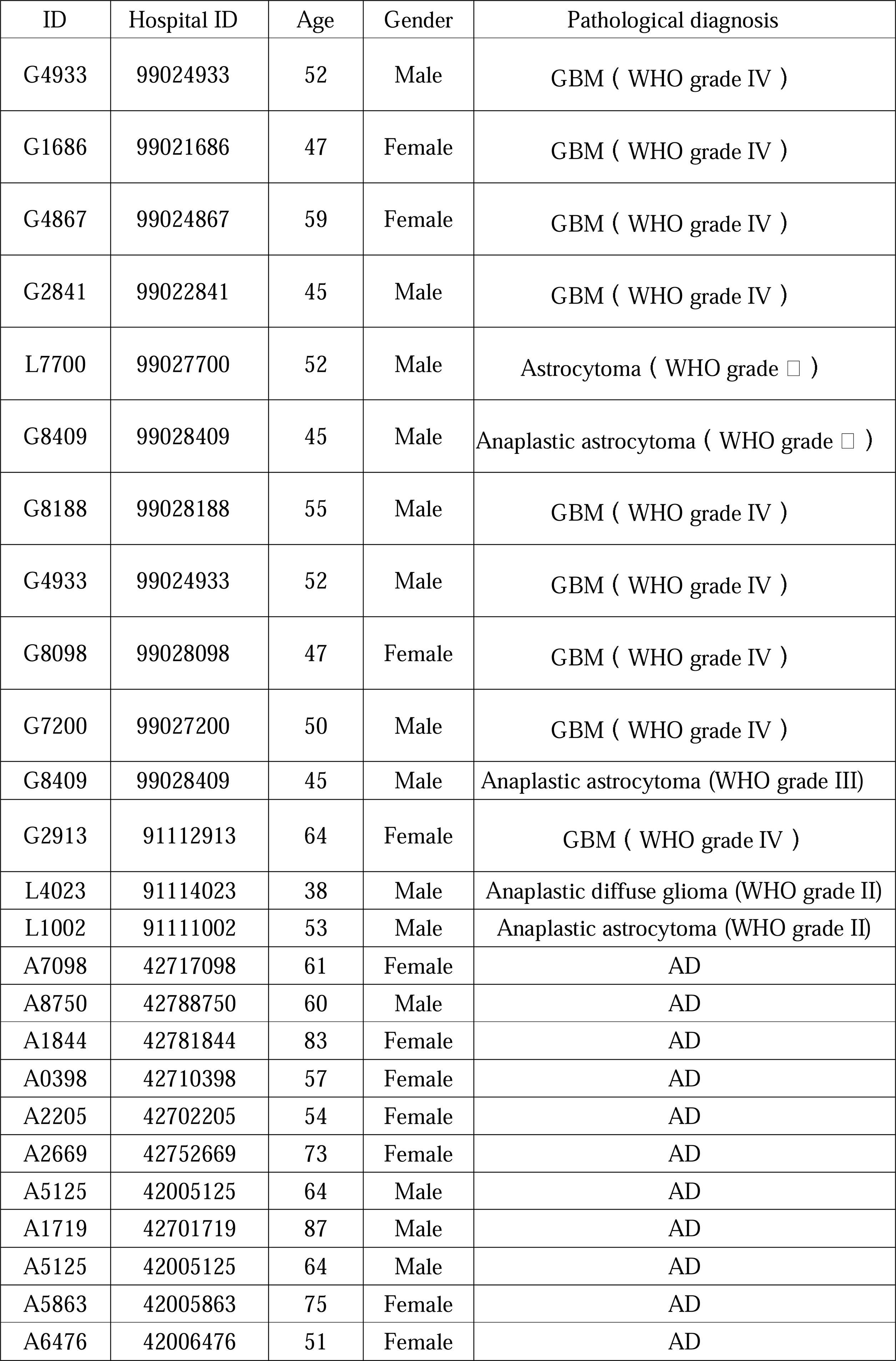
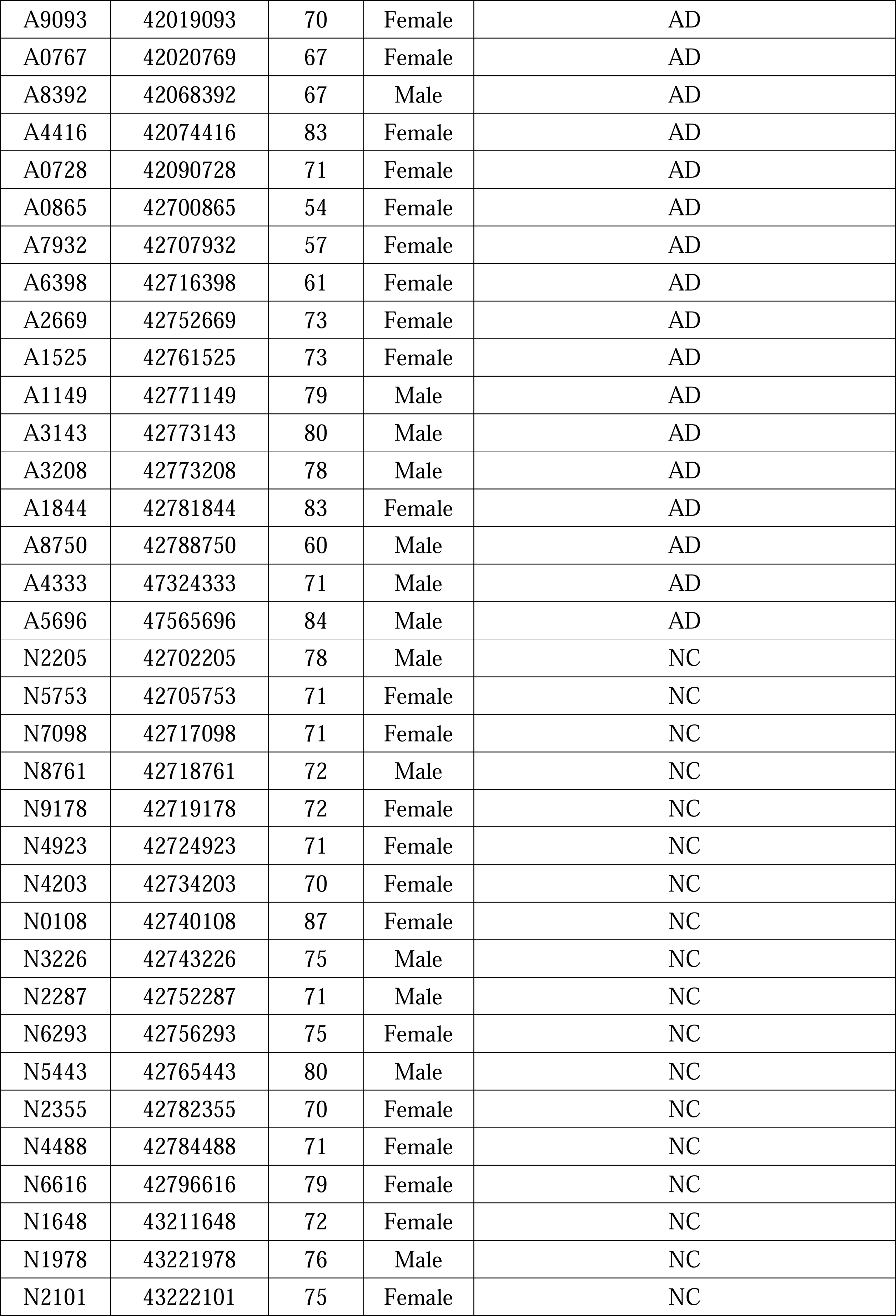
Clinical information.

| ID | Hospital ID | Age | Gender | Pathological diagnosis |
| --- | --- | --- | --- | --- |
| G4933 | 99024933 | 52 | Male | GBM ( WHO grade IV ) |
| G1686 | 99021686 | 47 | Female | GBM ( WHO grade IV ) |
| G4867 | 99024867 | 59 | Female | GBM ( WHO grade IV ) |
| G2841 | 99022841 | 45 | Male | GBM ( WHO grade IV ) |
| L7700 | 99027700 | 52 | Male | Astrocytoma ( WHO grade □ ) |
| G8409 | 99028409 | 45 | Male | Anaplastic astrocytoma ( WHO grade □ ) |
| G8188 | 99028188 | 55 | Male | GBM ( WHO grade IV ) |
| G4933 | 99024933 | 52 | Male | GBM ( WHO grade IV ) |
| G8098 | 99028098 | 47 | Female | GBM ( WHO grade IV ) |
| G7200 | 99027200 | 50 | Male | GBM ( WHO grade IV ) |
| G8409 | 99028409 | 45 | Male | Anaplastic astrocytoma (WHO grade III) |
| G2913 | 91112913 | 64 | Female | GBM ( WHO grade IV ) |
| L4023 | 91114023 | 38 | Male | Anaplastic diffuse glioma (WHO grade II) |
| L1002 | 91111002 | 53 | Male | Anaplastic astrocytoma (WHO grade II) |
| A7098 | 42717098 | 61 | Female | AD |
| A8750 | 42788750 | 60 | Male | AD |
| A1844 | 42781844 | 83 | Female | AD |
| A0398 | 42710398 | 57 | Female | AD |
| A2205 | 42702205 | 54 | Female | AD |
| A2669 | 42752669 | 73 | Female | AD |
| A5125 | 42005125 | 64 | Male | AD |
| A1719 | 42701719 | 87 | Male | AD |
| A5125 | 42005125 | 64 | Male | AD |
| A5863 | 42005863 | 75 | Female | AD |
| A6476 | 42006476 | 51 | Female | AD |
| A9093 | 42019093 | 70 | Female | AD |
| A0767 | 42020769 | 67 | Female | AD |
| A8392 | 42068392 | 67 | Male | AD |
| A4416 | 42074416 | 83 | Female | AD |
| A0728 | 42090728 | 71 | Female | AD |
| A0865 | 42700865 | 54 | Female | AD |
| A7932 | 42707932 | 57 | Female | AD |
| A6398 | 42716398 | 61 | Female | AD |
| A2669 | 42752669 | 73 | Female | AD |
| A1525 | 42761525 | 73 | Female | AD |
| A1149 | 42771149 | 79 | Male | AD |
| A3143 | 42773143 | 80 | Male | AD |
| A3208 | 42773208 | 78 | Male | AD |
| A1844 | 42781844 | 83 | Female | AD |
| A8750 | 42788750 | 60 | Male | AD |
| A4333 | 47324333 | 71 | Male | AD |
| A5696 | 47565696 | 84 | Male | AD |
| N2205 | 42702205 | 78 | Male | NC |
| N5753 | 42705753 | 71 | Female | NC |
| N7098 | 42717098 | 71 | Female | NC |
| N8761 | 42718761 | 72 | Male | NC |
| N9178 | 42719178 | 72 | Female | NC |
| N4923 | 42724923 | 71 | Female | NC |
| N4203 | 42734203 | 70 | Female | NC |
| N0108 | 42740108 | 87 | Female | NC |
| N3226 | 42743226 | 75 | Male | NC |
| N2287 | 42752287 | 71 | Male | NC |
| N6293 | 42756293 | 75 | Female | NC |
| N5443 | 42765443 | 80 | Male | NC |
| N2355 | 42782355 | 70 | Female | NC |
| N4488 | 42784488 | 71 | Female | NC |
| N6616 | 42796616 | 79 | Female | NC |
| N1648 | 43211648 | 72 | Female | NC |
| N1978 | 43221978 | 76 | Male | NC |
| N2101 | 43222101 | 75 | Female | NC |

